# Machine learning of brain-specific biomarkers from EEG

**DOI:** 10.1101/2023.12.15.571864

**Authors:** Philipp Bomatter, Joseph Paillard, Pilar Garces, Jörg Hipp, Denis Engemann

## Abstract

Electroencephalography (EEG) has a long history as a clinical tool to study brain function, and its potential to derive biomarkers for various applications is far from exhausted. Machine learning (ML) can guide future innovation by harnessing the wealth of complex EEG signals to isolate relevant brain activity. Yet, ML studies in EEG tend to ignore physiological artifacts, which may cause problems for deriving biomarkers specific to the central nervous system (CNS). We present a framework for conceptualizing machine learning from CNS versus peripheral signals measured with EEG. A common signal representation across the frequency spectrum based on Morlet wavelets allowed us to define traditional brain activity features (e.g. log power) and alternative inputs used by state-of-the-art ML approaches (covariance matrices). Using more than 2600 EEG recordings from large public databases (TUAB, TDBRAIN), we studied the impact of peripheral signals and artifact removal techniques on ML models in exemplary age and sex prediction analyses. Across benchmarks, basic artifact rejection improved model performance whereas further removal of peripheral signals using ICA decreased performance. Our analyses revealed that peripheral signals enable age and sex prediction. However, they explained only a fraction of the performance provided by brain signals. We show that brain signals and body signals, both reflected in the EEG, allow for prediction of personal characteristics. While these results may depend on specific prediction problems, our work suggests that great care is needed to separate these signals when the goal is to develop CNS-specific biomarkers using ML.

## Introduction

Electroencephalography (EEG) has a long history as a non-invasive technique for measuring brain activity in clinical research and practice. In the past decades, EEG has become increasingly popular as a technique for studying brain function in neurology (Gaubert et al., 2019; Jovicich et al., 2019; Schumacher et al., 2020; Sidorov et al., 2017; Sun et al., 2018; Zijlmans et al., 2012) psychiatry (Hegerl et al., 2012; Lenartowicz and Loo, 2014) and drug development (Janz et al., 2022; Leiser et al., 2011). EEG measures electrical potentials induced by cortical large-scale synchrony at dozens to hundreds of electrode locations on the scalp (Nunez and Srinivasan, 2006) and at temporal scales on the order of milliseconds to minutes (Buzsáki and Draguhn, 2004). The resulting multi-dimensional time series contain rich information about brain activity that can be quantified, e.g. as spectral power, spatial patterns, and waveform morphology (Jackson et al., 2019). Therefore, EEG is a valuable source of information that shows promise for deriving biomarkers of cognitive function, CNS pathology, and pharmacodynamics.

EEG signals are intrinsically complex. Analyses focusing on select frequencies, electrodes or time points can be successful in clinical settings characterized by global changes in EEG signals, e.g. related to changes in wakefulness or consciousness induced by sleep (Benca et al., 1999), severe brain injuries (Engemann et al., 2018; Schiff et al., 2014) or anesthesia (Purdon et al., 2013). More refined modeling, on the other hand, could uncover subtler EEG signatures and broaden the application of EEG. Propelled by advances in computer science, signal processing and the increasing availability of large EEG datasets, machine learning (ML) has emerged as a promising technology for isolating hidden patterns from complex EEG signals. ML, therefore, has the potential to unlock novel types of EEG biomarkers, e.g. to predict progression risk in neurodegenerative disorders (García-Pretelt et al., 2022; Gaubert et al., 2021) or to predict treatment success (Wu et al., 2020; Zhdanov et al., 2020).

As ML methods, including deep learning (DL), for EEG are rapidly developing, the field has not yet converged on methodological standards and best practices (Roy et al., 2019). This increases the researcher’s degrees of freedom, and consequently the variability of results, which may hamper unlocking the potential of ML for improving EEG analysis. One potentially important source of such variability arises from handling artifactual signals in the EEG. A recent systematic review of the ML & DL literature for EEG (Roy et al., 2019), found that the majority of studies (72%) did not perform any explicit removal of physiological artifacts related to peripheral body signals (e.g. eye blinks, muscle & cardiac activity) that are well known to leak into the EEG and can overshadow the brain signal of interest. This is particularly problematic as these peripheral sources are often modified by medical conditions and other individual factors and might therefore be predictive too (Golding et al., 2006; Lage et al., 2020; Lindow et al., 2023), which is entirely ignored if EEG is left unprocessed as was recently advocated for (Delorme, 2023). Depending on the relationship between artifacts and variables of interest, high-capacity ML techniques such as deep neural networks may automatically learn how to filter out artifact-generating sources as irrelevant noise, or, instead will use non-brain information to minimize prediction error (Jochmann et al., 2023). The latter case would hamper unambiguous interpretation as brain-specific biomarkers.

Artifact removal techniques such as independent component analysis (ICA) or signal-space projection (SSP) have proven effective in reducing the leakage of signals from non-brain generators into the EEG (Hyvärinen et al., 2004; Uusitalo and Ilmoniemi, 1997). In recent years, technological advancements have made it easier to automate and scale these artifact removal techniques (Jas et al., 2017; Pion-Tonachini et al., 2019; Zhang et al., 2021). Yet, they have not been systematically studied in the context of machine learning pipelines for biomarker learning. It remains unclear if and how artifact removal procedures that isolate brain signals from body signals impact model performance and to what extent ML models make use of EEG signals induced by peripheral physiological generators.

This work had two principal scientific objectives. 1) We aimed at formalizing the importance of considering potentially predictive physiological signals in a conceptual framework for building interpretable brain-specific EEG biomarkers with ML. 2) We tested if ML models make systematic use of peripheral non-brain signals if EEG signals are not sufficiently preprocessed.

We focused on ML approaches for subject-level prediction, where one data point contains one EEG recording and one single outcome measure (Fruehwirt et al., 2017; Sabbagh et al., 2020; Wu et al., 2020) as compared to event-related modeling in cognitive decoding (King and Dehaene, 2014; Stokes et al., 2015) or brain-computer interfaces (BCI) (Abiri et al., 2019; Congedo et al., 2017). As ML requires training data, age and sex prediction are promising example problems that are readily accessible across data resources and have received increasing attention in human neuroscience. For example, the brain age approach (Cole et al., 2019; Smith et al., 2019) encapsulates patterns of brain aging via age prediction models, which, evaluated on atypical or clinical populations, can show informative biases like over-prediction of the chronological age (Cole et al., 2018; Denissen et al., 2022). A similar idea has been recently explored for sex prediction (Floris et al., 2023). Both age and sex prediction are actively investigated with EEG (Binnie et al., 2021; Engemann et al., 2022; Jochmann et al., 2023; Sun et al., 2019; van Putten et al., 2018). To study the interplay between predictive brain and body signals captured by the EEG through age and sex prediction, we chose publicly accessible datasets with wide age ranges for which at least 1000 data points were available, i.e. TDBRAIN, the Two Decades of Brainclinics (van Dijk et al., 2022), and TUAB, the Temple University Hospital Abnormal Corpus (Obeid and Picone, 2016).

We used a generic framework suited for expressing prior biological assumptions. Previous work has obtained promising results for subject-level prediction of age from EEG by relying on the between-electrodes covariance matrices from different frequency bands as model inputs (Engemann et al., 2022). This approach is backed by statistical theory and defines mathematical tools from Riemannian geometry for building prediction algorithms on covariance manifolds (Barachant et al., 2010; Congedo et al., 2017; Sabbagh et al., 2020). The resulting models are effective at suppressing the distorting effects of volume conduction and electrical potential spread (Sabbagh et al., 2020). Our theoretical framework follows this line of research and extends it by making the role of non-brain predictive artifacts explicit. Moreover, our approach is committed to avoiding hand-picking of frequencies or band definitions and to improving interpretability against classical spectral measures. We therefore computed the covariances using complex Morlet wavelets (Morlet et al., 1982) that have a long tradition in EEG signal analysis (Cohen, 2019; Hipp et al., 2012; Tallon-Baudry et al., 1996). This allowed us to cover the entire frequency spectrum with fine-grained resolution. As a second complementary generic approach, we benchmarked a convolutional neural network (Schirrmeister et al., 2017) that has the capability to learn custom oscillatory motifs.

## Results

### Conceptual framework for building brain-specific prediction models with EEG

We first developed our methodological framework (Figure 1). Prior work (Sabbagh et al., 2020) proposed a generative modeling framework for regressing biomedical outcomes on cortical activity in the presence of volume conduction. A central feature of that work are statistical guarantees that yield unbiased prediction models without approximation error – also termed statistical consistency – given specific model assumptions of, e.g. linear field spread, and a log linear relationship between brain activity and the outcome (eqs. 2 and 3). Moreover, the framework can accommodate matrix-rank deficiencies caused by artifact cleaning (Absil et al., 2009; Sabbagh et al., 2019), which is of central importance in our context.

**Figure 1.**
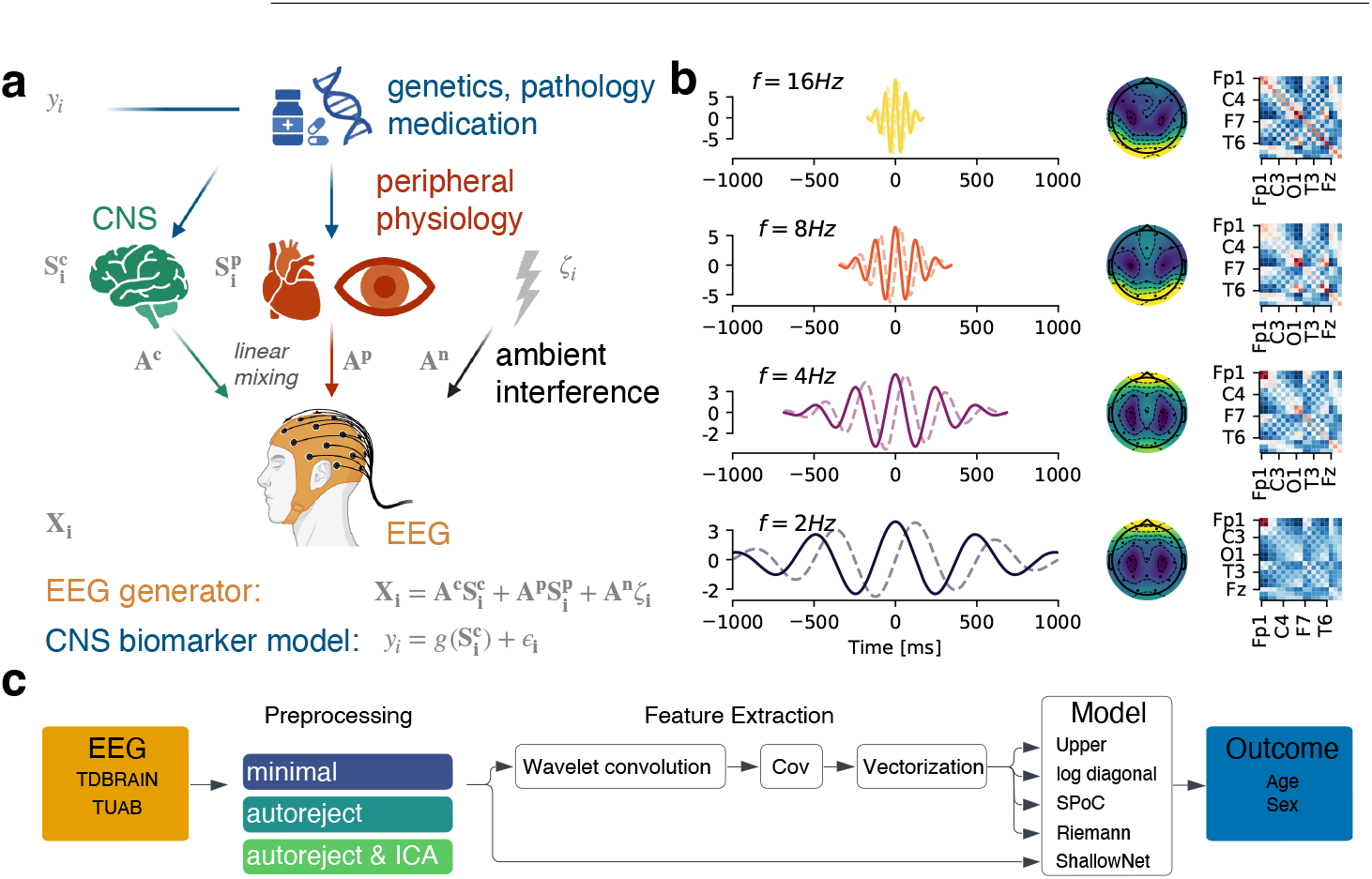
Conceptual framework for biomarker learning from EEG. Conceptual framework for biomarker learning from EEG. (**a**) Individual genetics, medical conditions, and interventions among other factors affect the brain (CNS) and the body (periphery) simultaneously, enabling predictive modeling as outcome measures related to these conditions become correlated to physiological measures such as EEG. Predictive CNS generators, predictive peripheral generators (e.g., ocular, cardiac, muscular) and non-predictive ambient noise (electronics, loose electrodes, electromagnetic interference) induce EEG signals through linear mixing. An ideal CNS biomarker isolates brain-related signals and ignores the other generators contributing to the EEG signal. (**b**) Families of complex Morlet wavelets (left) offer useful representations for disentangling signal generators inducing EEG patterns in different frequencies (solid and dotted lines represent real and imaginary parts of the complex valued kernel). Beyond classical log power topographies (middle), we derived covariance matrices (right) through wavelet convolutions. Covariances play a central role in machine learning (ML) algorithms for EEG as they define representations that help mitigate distortions and biases due to linear source mixing. (**c**) Data processing and predictive modeling pipeline. To study the impact of different EEG generators on ML pipelines, we varied the preprocessing (*minimal* - numerically stabilizing processing, *autoreject* - removal of ambient noise and high-amplitude signals, *autoreject & ICA* – additional removal of peripheral artifacts). We focused on Morlet wavelets and covariance-based approaches that emerged as informative baselines in previous work as they imply different underlying hypotheses about the regression function and make different use of spatial information (Table 1). We also investigated a convolutional neural network (*ShallowNet*) capable of learning custom, temporal filters from the data, which leads to increased model capacity and enables learning of features sensitive to changes in the shape of oscillations or bursting events. **Figure 1 – Figure supplement 1**: Prediction results with wavelets versus conventionally defined frequency bands (IPEG).

Here, we extended that framework to explicitly reflect the role of potentially predictive body signals as compared to non-predictive noise (Figure 1, ÿeq. 2). This inspired us to propose a stringent definition of CNS biomarkers for which we require that the EEG model isolates CNS components from noise and peripheral signals (eqs. 2 and 3). At the theoretical level, this formulation allowed us to see that statistically consistent regression models for prediction from EEG brain activity Sabbagh et al. (2020, 2019) from EEG-sensor-space covariance matrices (eq. 4) are also consistent for predicting from non-brain activity leaking into the EEG, unless dedicated artifact cleaning is applied. It is therefore not necessarily a safe assumption that such machine learning models will automatically learn to ignore artifacts with enough training data (see formal analysis presented in methods, (eqs. 12 to 15). If artifacts are uncorrelated with the outcome of interest, one might choose to leave the data unprocessed and thereby potentially even increase the robustness of the learned representation. However, prior studies have shown that peripheral signals and EEG artifacts can be systematically modulated in different patient groups (Golding et al., 2006; Jongkees and Colzato, 2016; Lindow et al., 2023; Wilkinson and Nelson, 2021). This motivated us to systematically study the relationship between different EEG components attributed to CNS versus peripheral generators, which typically differ in terms of spectral and spatial patterns.

In support of this purpose, we combined ML with EEG representations based on Morlet wavelets (Figure 1a, eq. 4; Morlet et al. 1982; Tallon-Baudry et al. 1996). Our approach to spectral analysis chooses a log-frequency parametrization defining the wavelet families using a base-2 logarithmic grid (Hipp et al., 2012), which is tailored to the natural scaling of brain rhythms that are rather lognormal (Buzsáki and Mizuseki, 2014). That is, the spectral resolution is higher at lower frequencies, and accordingly the spectral smoothing is greater at higher frequencies. This representation is well established for implementing spectral EEG measures (e.g. log power, eq. 7) and a common choice in clinical biomarker studies (Frohlich et al., 2019; Hawellek et al., 2022; Janz et al., 2022). Beyond classical EEG metrics, here, we adapted the Wavelet approach to further derive advanced representations for state-of-the-art ML methods developed for EEG (Figure 1a,Figure 1b).

As our principal ML strategy, we focused on the family of covariance-based prediction models (Congedo et al., 2017; Grosse-Wentrup and Buss, 2008; Koles et al., 1990) that were theoretically and empirically analyzed in previous work (Sabbagh et al., 2020). These models provide useful baselines as they enjoy statistical guarantees under different assumptions, such that the comparison of their performance can hint at characteristics of the data-generating mechanism (cf. *Prediction algorithms*). So far, these models have been studied with conventional frequency bands and other than in previous work, we estimated frequency-specific covariances from the real part of the Morlet wavelet representation (Figure 1b). Additional validation confirmed that covariance-based models performed consistently better when implemented under the wavelet approach. For a comparison with frequency bands proposed by the International Pharmaco-EEG Society (IPEG Jobert et al. 2012), see also Figure 1 – Figure supplement 1.

We also explored a flexible convolutional neural network approach: *ShallowNet* (Schirrmeister et al., 2017) operates directly on EEG time series data and can overcome the potential limitations of the assumptions of sinusoidal brain oscillations at fixed frequencies (Jackson et al., 2019). Together, these models should be reasonably representative of state-of-the-art ML models used in EEG research.

We then used this combination of a high high-resolution spectral representation and powerful covariance-based ML models that can leverage fine-grained spatial features to investigate the impact of artifact preprocessing. We performed analyses across different levels of preprocessing, designed to vary the extent to which environmental versus physiological artifacts are cleaned (Figure 1c).

### Impact of Preprocessing on Model Performance

We applied our framework to investigate the contribution of brain and non-brain signals to the performance of the different ML models in two benchmark tests, age, and sex prediction. We compared model performance across different levels of preprocessing to assess if ML models would benefit, suffer or be unaffected by artifact removal. We compared the 10-fold cross-validation results for different models on either minimally, moderately, or extensively preprocessed input data (Figure 2, Table S1, Table S2). Minimally preprocessed data was filtered and resampled. Moderate preprocessing made additional use of *autoreject* (Jas et al., 2017), a method that discards and interpolates bad channels and segments (AR) to remove

**Figure 2.**
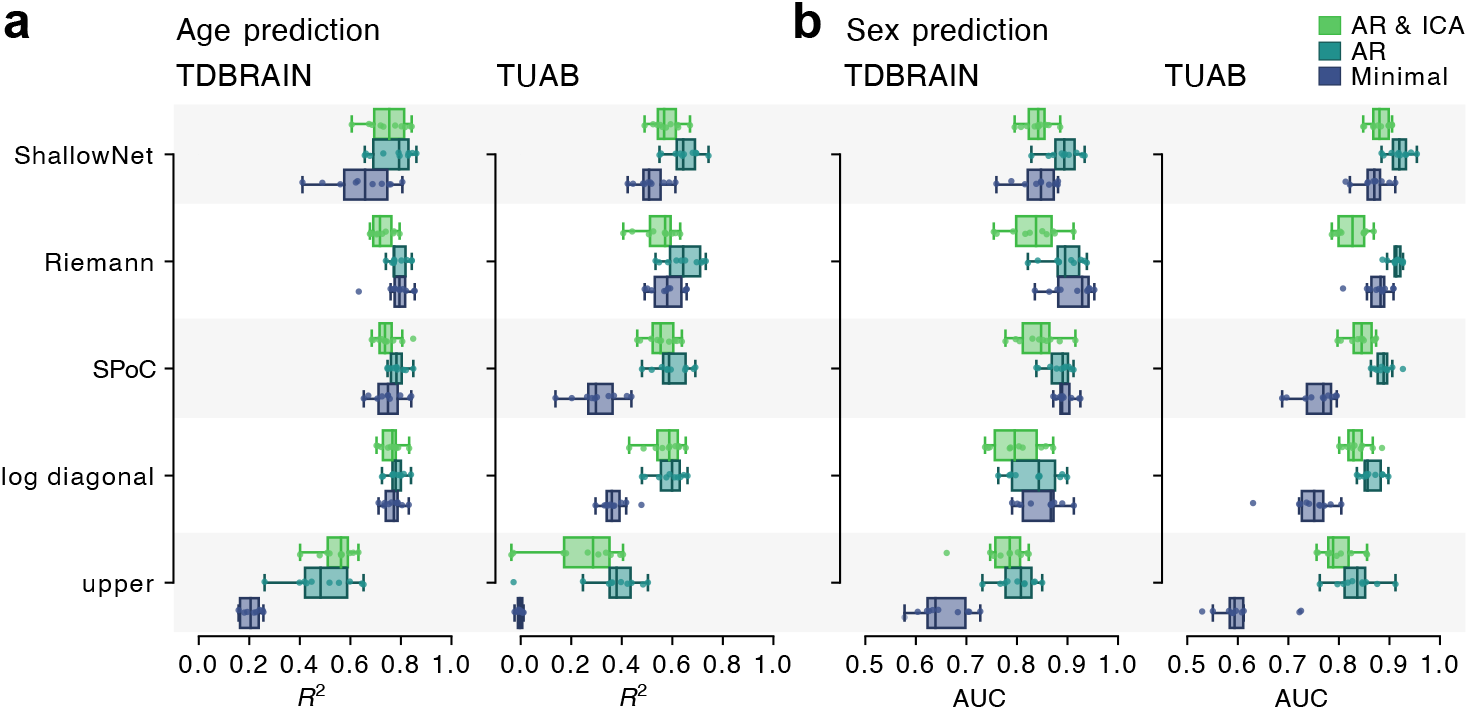
Impact of artifact removal on EEG prediction performance. (**a**): age prediction and (**b**): sex classification on the TDBRAIN and the TUAB datasets. Traditional boxplots show 10-fold-cross validation distributions (middle line represents the median). Color depicts the degree of EEG processing (*minimal, autoreject, autoreject & ICA*). Rows present model architectures: Ridge regression & classification based on covariances with upper vectorization (upper), the log of the variance components (*log diagonal*), supervised spatial filtering (*SPoC*), the Riemannian tangent space (*Riemann*) and a convolutional neural network operating on raw EEG time series (*ShallowNet*). For readability, the horizontal axis was clipped at *R*^2^ = − 0.1, occluding folds with prediction failure for the *upper* model. Removing noisy channels and high-amplitude data segments (*autoreject*) often led to improved performance. The Riemannian model benefited least from intense preprocessing, whereas simpler models (*upper, log diagonal*) could be substantially improved by preprocessing. Performing additional ICA-based rejection of artifacts (muscles, eye blinks, etc.) often lowered the prediction performance, suggesting that bodily, non-brain generators of the EEG can include predictive information.

**Figure 2 – Figure supplement 1**: Impact of processing on EEG power spectra, topographies and covariances (TDBRAIN).

**Figure 2 – Figure supplement 2**: Impact of processing on EEG power spectra, topographies and covariances (TUAB).

high-amplitude artifacts produced by ambient interference, electronics or loose electrodes. Extensive preprocessing added explicit removal of physiological artifacts using ICA (AR & ICA) supported by automatic labeling of brain and artifact components (Ablin et al., 2018; Hyvärinen and Oja, 2000; Pion-Tonachini et al., 2019).

The depth of preprocessing affected total power, relative power and covariances across frequencies (Figure 2 – Figure supplement 1 & Figure 2 – Figure supplement 2). Grouping of log power spectra by age showed differences that may enable prediction, regardless of preprocessing. Except for the *upper* model, all models showed above-chance prediction regardless of processing choices (Figure 2a,b). Pooling over models and datasets, we can observe that *autoreject* led to an improvement in *R*^2^ over minimally preprocessed data of 0.151 (CI_95%_ = [0.076, 0.236]) for age prediction (Figure 2) and in area under the curve (AUC) of 0.071 (CI_95%_ = [0.021, 0.126]) for sex prediction (Figure 2b). The additional ICA step did not yield further improvements and instead lowered the performance by on average -0.036 (CI_95%_ = [ −0.061, −0.004]) in *R*^2^ and -0.047 (CI_95%_ = [ −0.057, −0.037]) in AUC compared to *autoreject* preprocessing alone.

Detailed model comparisons revealed that simpler models tended to benefit more from preprocessing. The *upper* model (eq. 8), for instance, failed on the task of age prediction on TUAB when only *minimal* preprocessing was applied to the input, and performance was substantially improved with preprocessing. A similar effect was observed for the *log diagonal* and *SPoC* models. The *Riemann* model and the *ShallowNet* benefited less from preprocessing with *autoreject*. This might be explained by their increased capacity to use the wealth of information in raw signals or the covariance matrix to suppress irrelevant signals. All cross-validation results including additional metrics are printed in Table S1 and Table S2 to facilitate comparisons with other studies.

In summary, our results suggest that removal of high-amplitude artifacts using *autoreject* can be beneficial across models, whereas refined ICA-based removal of physiological artifacts might hamper predictive information.

### Exploring the relative contribution of peripheral non-brain signals

That ICA-based removal of non-brain signals (including muscle activity, eye movements or cardiac activity) led to lower prediction performance could imply that these non-brain signals contained information about the outcome. We next performed an in-depth investigation of the predictive value of these signals that are typically treated as artifacts. To approximate a decomposition of the model performance into brain and non-brain contributions (eqs. 12 to 15), we explored two complementary approaches: (1) reconstruction of EEG features from brain and artifactual ICs, and (2) computation of EEG features from auxiliary channels, where available. Specifically, we focused on the auxiliary channels used to record ocular, muscular, and cardiac activity in the TDBRAIN dataset (see *Approximate subspace regression and classification*).

If the drop in performance after ICA preprocessing is due to the removal of predictive information, we should achieve above-chance prediction performance when using only the removed signal as input. To investigate this hypothesis, we first used ICA on the TUAB dataset where no auxiliary channels were available (Figure 3). We reconstructed the signal from the subspaces spanned by the rejected ICA components. As the ICLabel algorithm (Pion-Tonachini et al., 2019) used for labeling of independent components provides categories for the rejected components, we also reconstructed the signal for particular classes of artifacts (e.g. ocular or muscle artifacts).

**Figure 3.**
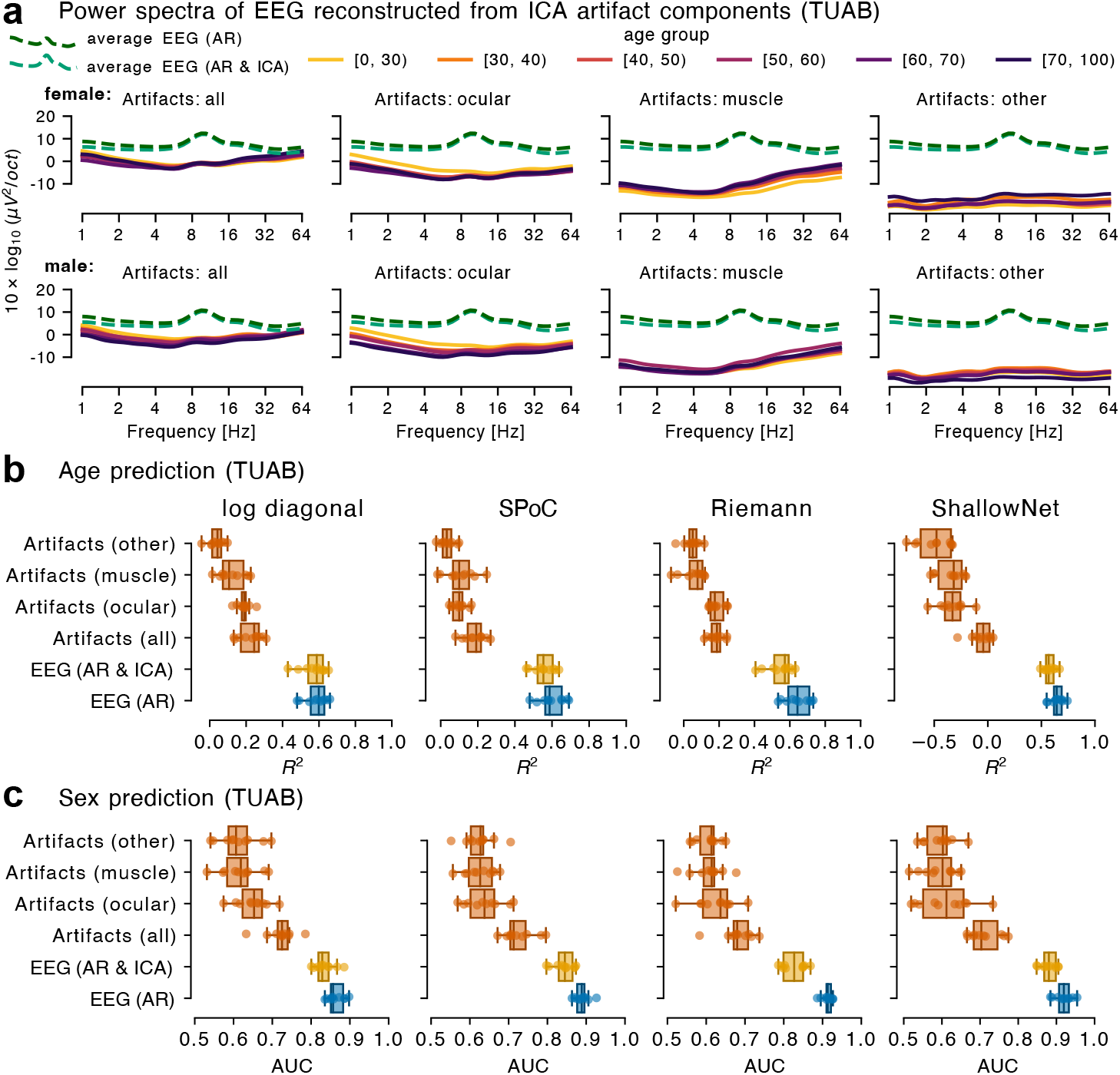
Exploration of the contribution of CNS and peripheral signals as extracted from ICA. **(a)** Power spectra of EEG reconstructed from ICA components identified as reflecting brain signals and artifacts, averaged across electrodes. Power in ocular artifact subspaces is concentrated at low frequencies and muscle activity is concentrated at high frequencies. Note that the overall lower amount of power for artifacts compared to the cleaned signal is partially explained by the averaging across all electrodes: artifacts are typically concentrated in a subset of the electrodes. Importantly, alpha power (8-12 Hz) was largely preserved in EEG and virtually absent in ICA-reconstructed artifact signals. **(b)** and **(c)** show model comparisons for cleaned EEG versus ICA-reconstructed artifact signals. Performance was higher after cleaning with AR (blue) and AR & ICA (yellow), yet the ICA reconstructions of artifact subspaces (orange) also contained predictive information. The highest performance was achieved after AR preprocessing, which eliminated large artifact sections but did not eliminate the contribution of physiological artifacts.

Inspecting the power spectra of the reconstructed artifact signals, ocular artifacts were dominated by low-frequency power, whereas muscle artifacts contained mostly high frequency power (Figure 3a). Moreover, the alpha-band peak above 8 Hz was preserved after ICA-cleaning of the EEG (signal-subspace) and not present on the artifact-subspace power spectra (Figure 3a). Grouping by age and sex on subspace power spectra showed average power differences that may enable prediction. We used 10-fold cross-validation to gauge model performances for age and sex prediction (Figure 3b,c). The subspace models achieved above-chance performance, suggesting that non-brain signal generators are predictive of age and sex. The only exception to this pattern was the *ShallowNet* performance for age prediction, potentially related to data requirements of complex regression models in situations with low signal-to-noise ratio. Jointly predicting from all artifact sources together yielded better performance than predicting from any individual artifact class, but performance results for individual artifact classes were still above chance. Importantly, the performance of models using clean EEG were substantially better and performance distributions were non-overlapping with the artifact subspace models.

The second approach, using auxiliary channels instead of ICA subspaces, was explored on the TDBRAIN dataset (see *TDBRAIN dataset*). The horizontal and vertical EOG electrodes are placed close to the eyes and thus primarily pick up eye movements, which is reflected in high power at low frequencies. The ECG electrode is placed at the cervical bone and provides a measurement of cardiac activity. Finally, the EMG electrode is placed on the right masseter muscle and measures (jaw) muscle activity, reflected in large amounts of high-frequency power (Figure 4a). As we expected, comparing power spectra, all auxiliary channels appeared markedly different from the EEG channels. However, we can still discern alpha peaks between 8 and 12 Hz in the EOG and EMG channels, suggesting that auxiliary channels also picked up brain activity to some extent. As before, the grouping by age and sex on the auxiliary-channel power spectra reveal differences that may enable prediction. In line with results from the ICA-based analysis, predictions based on auxiliary channels were substantially better than chance (Figure 4b,c). In many cases, data from EOG channels explained the larger part of the performance obtained with auxiliary channels.

**Figure 4.**
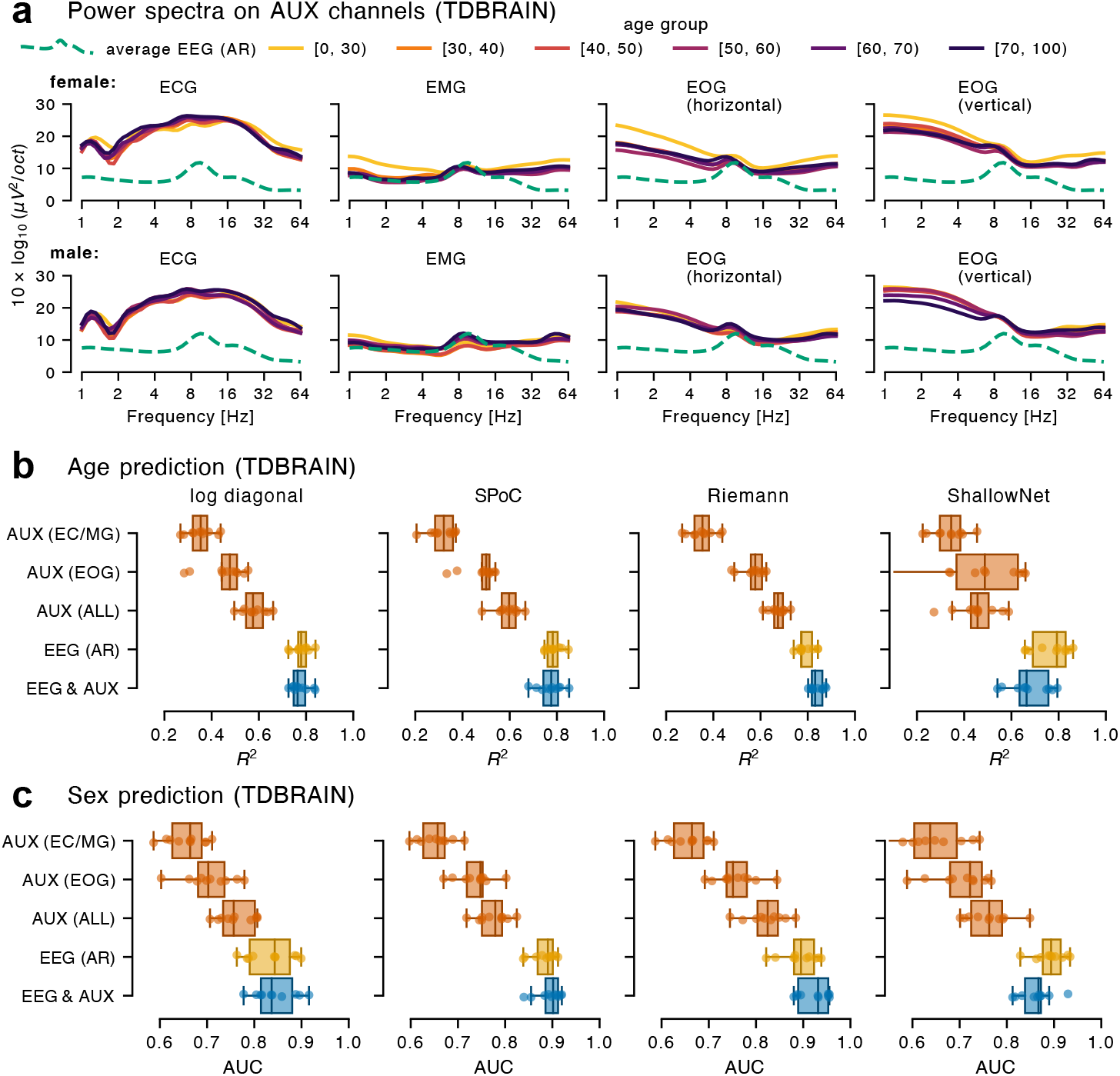
Exploration of the contribution of brain and body signals through analysis of auxiliary (AUX) channels. **(a)** AUX channel power spectra generally exhibit different characteristics than EEG (cleaned with *autoreject*, averaged across electrodes). **(b)** and **(c)** show comparisons of model performance with EEG versus auxiliary physiological channels (AUX) for age and sex prediction, respectively. It can be seen that modeling from AUX inputs (orange) leads to systematic prediction performance across tasks and model architectures. The effect of AUX inputs related to ocular activity explained much of the performance of all AUX inputs together. Predicting from EEG inputs (yellow) yielded consistently higher performance. As a tendency, model architectures that achieve better results for EEG inputs also achieve better results for AUX inputs. Combining EEG and AUX features (blue) did not result in systematically better performance. These results suggest that the predictive information conveyed by AUX channels is already captured by the EEG channels.

Yet, the combination of all available auxiliary channels led to the highest performance compared to individual auxiliary channels. Again, the performance achieved with EEG data was substantially higher, as evidenced by non-overlapping or weakly overlapping cross-validation distributions. Strikingly, despite the performance observed with auxiliary channels, combining EEG and auxiliary channels did not lead to consistent improvements over pure EEG data for most models. This suggests that the predictive information present in the auxiliary channels was already present in the EEG signal and explained part of its performance.

In sum, our findings suggest that across datasets and tasks, non-brain signals contain information predictive of age and sex but that the main driver of prediction performance is brain related.

### Model exploration through spectral profiling of prediction performance

The wavelet-based framework developed in this work not only offers competitive prediction performance (Figure 1 – Figure supplement 1). It provides additional opportunities for model interpretation. How individual frequencies contribute to model prediction can provide insights about the underlying physiological processes. E.g. in motor tasks, researchers have been interested in specific frequency ranges (beta frequency range), possibly reflecting strong associations of outcomes with oscillatory activity in the motor cortex (Peterson et al., 2019; Schoffelen et al., 2011). On the other hand, specific frequency ranges might not play a prominent role if predictive brain sources are widely spread across cortical networks. Moreover, structural anatomical characteristics may systematically influence propagation of brain activity, as recently hypothesized for sex prediction from EEG (Jochmann et al., 2023). Also, changes of states of consciousness (awake, sleep, coma) that go hand in hand with global changes of the EEG signal may enable decoding from broadband power. This included certain drugs like anesthetics (Bojak and Liley, 2005; Drummond et al., 1991).

Comparing models with a restricted frequency range versus all frequencies, allowed us to explore the nature of the predictive signal. We focused this analysis on the *SPoC* model, which strikes a good compromise between computation time and model performance. As covariances at a single frequency cannot capture local changes across frequencies, we built models capable of deriving local contrasts between neighboring frequencies (Figure 5, left subpanels, blue lines). For this purpose, we extracted covariances from 5 neighboring wavelets around the center frequency *f* spanning one octave (see *Background: Spectral analysis and machine learning for EEG biomarkers*). In addition, we averaged covariances (with fixed equal weights) across the same neighboring frequencies (Figure 5, left subpanels, yellow lines). Comparing these two approaches allowed exploring the complexity of local information as performance should be equal if local changes in the spatial patterns along the spectrum contain no information.

**Figure 5.**
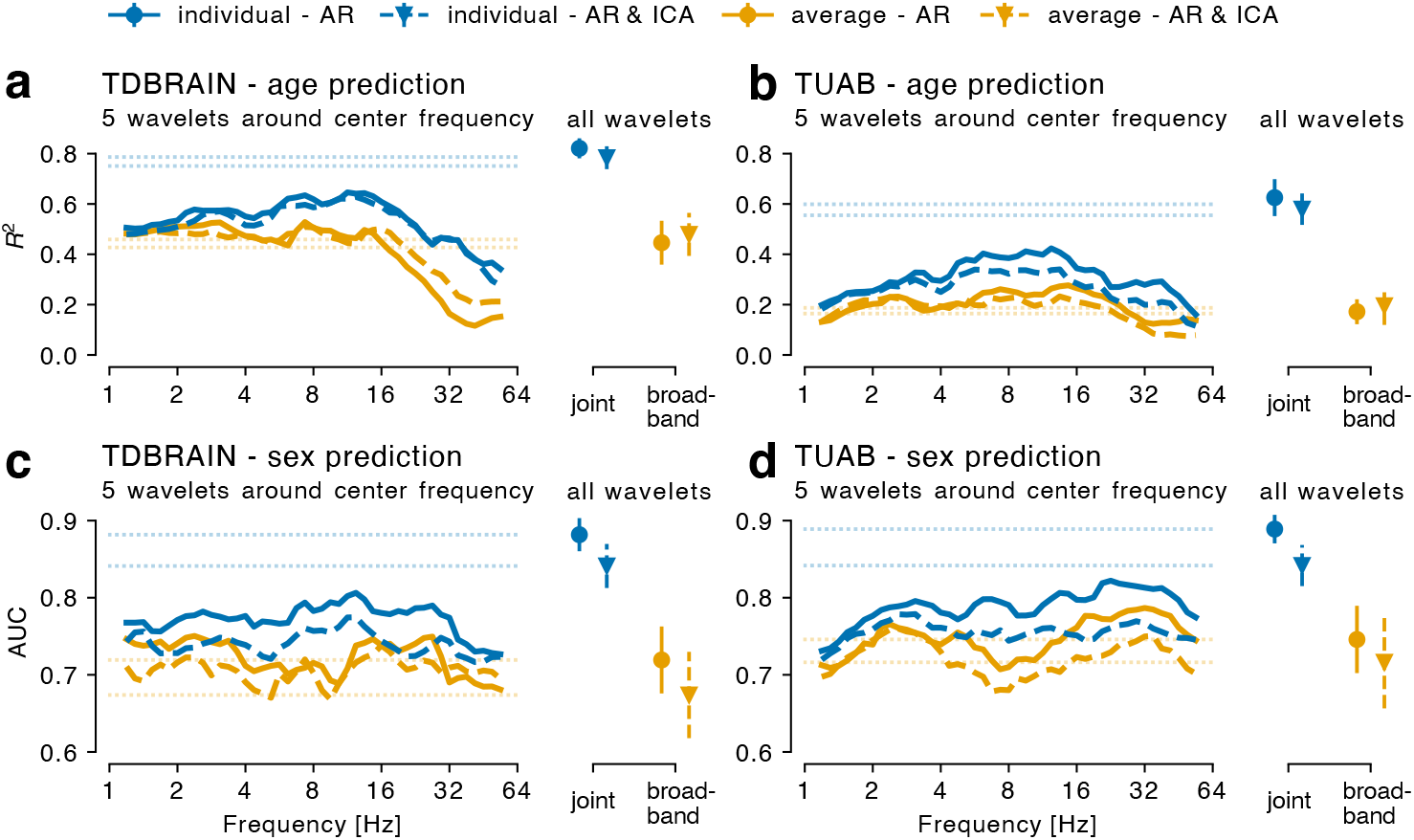
Spectral profiling of predictive EEG signatures using the wavelet framework. We inspected the *SPoC* pipeline on TDBRAIN (**a, c**) and TUAB (**b, d**) datasets for age (**a, b**) and sex (**c, d**) prediction by confronting frequency-wise models (left subpanels) with full cross-frequency models (right subpanels, all wavelets). For frequency-wise modeling, we used 5 wavelets around the center frequency (x-axis), from which covariances were averaged (yellow) or individually combined by the model (blue). Solid versus dashed curves depict the preprocessing, i.e., *autoreject* only and additional ICA, respectively. Horizontal lines provide an orientation with regard to the results based on all wavelets (right subpanels). The joint model with all individual wavelets represents the previously investigated standard models. The broadband model averaged all covariances across all frequencies to capture the global signal. It can be seen that letting the model combine the frequencies consistently led to better results, pointing at spectrally distributed information.

To put performance based on these models with a restricted frequency range around center frequencies into perspective with previous results, we replotted the full model joining all frequencies as input (Figure 5, right subpanels, blue markers), corresponding to results shown for *SPoC* in Figure 2. In addition, we conducted a control analysis using the average of all covariances as input, which basically corresponds to the covariance of the broad-band signal that lacks explicit frequency information (Figure 5, right subpanels, yellow markers). Furthermore, we explored the effect of preprocessing on performance in local frequency regions as compared to global effects (Figure 5, dotted lines and markers).

We first focussed on age prediction from EEG cleaned with *autoreject* (solid lines). For TDBRAIN (Figure 5a), performance peaked between 8-16 Hz, around *R*^2^ scores of 0.6. A similar peak emerged below 8 Hz. By comparison, adaptively joining all wavelets (as in our previous analysis), increased the *R*^2^ score to around 0.8. A similar picture emerged for TUAB (Figure 5 5b), with *R*^2^ peaking around 0.4 at similar frequencies, whereas, joining all wavelets increased the score to around 0.6. For orientation, uncertainty estimates (SD) are displayed on the right subpanels. This suggests that the information from different frequencies was complementary for age prediction on both datasets. Applying averaging (yellow lines), a clear drop in performance of around 0.2 in terms of R2 scores was observed on, both, local-frequency (Figure 5a,b, left subpanels) and global broadband models (Figure 5a,b, right subpanels), pointing at complex and local spectral information that is lost upon averaging.

A similar picture emerged for sex prediction (Figure 5c,d). For TDBRAIN, peak AUC scores of about 0.8 were observed between 8 and 16 Hz (Figure 5c). Joining all frequencies lifted the AUC score close to 0.9. This was highly similar for TUAB (Figure 5d), except that peak performance was observed above 16 Hz. By comparison, averaging (yellow lines) decreased the performance to AUC scores closer to 0.7.

To connect this analysis with our previous results, we analyzed changes in performance profiles as further artifact removal with ICA was added (dashed lines).

Changes in performance were small for age prediction in general, especially for the TDBRAIN dataset where frequency-specific models seemed unaffected. Some reduction in prediction performance at the peak above 8 Hz from 0.4 to around 0.3 is seen on the TUAB dataset. By comparison, the effect of removing artifacts was more visible for sex prediction, also for global combined models.

Taken together, spectral profiling revealed some frequency specificity in prediction performance while prediction was broadly possible across all frequencies. Models combining frequencies at local or global ranges always clearly outperformed averaged models, pointing at synergistic information across the frequency spectrum and ruling out trivial offsets or broadband effects as main drivers of the prediction.

## Discussion

Over the past decade, important advances have been made in EEG-based biomarker exploration with ML. To fully harness the potential of ML for EEG biomarkers, it will be important to optimally use neuroscientific and biophysiological insights from EEG research. Incorporating such prior knowledge into machine learning models can endow them with theoretical grounding and increase their robustness in wide-data regimes (few training data points, many variables) dominating human neuroscience. Our literature review identified an important gap in the conceptualization and practical handling of signal contributions to the EEG originating from non-brain, peripheral generators. Peripheral signals are rigorously treated as artifacts in classical EEG methods but are commonly ignored in ML work. This is potentially due to widespread enthusiasm about the capability of ML models to detect the hidden patterns of interest in data and ignore the noise.

Our work addresses the need for a practical yet theoretically grounded machine learning methodology for biomarker discovery and development with EEG. Our proposed framework carefully reconsidered previous theoretical results on regression models for predicting from brain activity in the presence of field spread and volume conduction. As biomedical conditions and therapeutic interventions can affect, both, the brain and the body, related outcomes can potentially be predicted from, both, brain and body signals. This insight has inspired us to study state-of-the-art ML algorithms in empirical benchmarks designed to evaluate and isolate the differential contributions of brain versus body signals mixed in the EEG. Our benchmarks demonstrate that exemplary age and sex prediction problems were substantially affected by non-brain signal generators if these were not explicitly handled.

A key insight obtained from our conceptual analysis and our empirical work is that prediction from EEG signals will only yield a brain-specific biomarker model if bodily signals are explicitly removed or controlled for.

Furthermore, our empirical results provide new insights into the prediction algorithms, extend the scope from age prediction to sex classification, and unlock model interpretation techniques through the proposed wavelet framework. In the following sections, we provide deep dives into some of the details and their practical implications.

### Artifact removal is essential for learning interpretable CNS biomarkers

The wavelet methodology allowed us to compile new benchmarks, visualization and analysis techniques for studying the interplay between CNS and peripheral EEG generators. Our benchmarks on artifact removal revealed a colorful picture. For all ML models, removal of high-amplitude artifacts via *autoreject* improved prediction performance (Figure 2), which contrasts a widespread view according to which *minimal* processing of EEG is preferable for ML (Delorme, 2023; Roy et al., 2019). On the other hand, additionally regressing out artifacts related to peripheral signals with ICA consistently led to a decrease rather than a further improvement in performance, which suggests that there is predictive information in these signals.

One common argument presented in the context of model interpretation is that one cannot tell ad-hoc whether certain features are necessarily bad because the model might use them to denoise the predictive function (Haufe et al., 2014). For instance, if the model has access to a channel that strongly reflects ocular artifacts, it could, in principle, use this channel as an artifact indicator and down-weight the importance of artifacted segments. This might motivate researchers to keep the data minimally processed (Roy et al., 2019). We see our theoretical framework and empirical results in disagreement with this view. The key point, formalized in our generative model, is that it matters whether these artifact signals are themselves correlated with the outcome. If one expects artifact signals to be useful— not because of their own predictive value but only as a means for the model to denoise the actual signal of interest–– one should also expect the removal of high-amplitude artifacts, related to bad channels and segments, to lead to a decrease in model performance. Our benchmarks show the opposite. It is thus much more plausible that the decrease in performance after removal of physiological artifacts with ICA is not explained by the prediction model’s limited capacity to denoise the signal, but instead because the removed signals are themselves predictive of the outcome.

A related argument would be to motivate the omission of preprocessing based on the emerging literature on data augmentation in EEG, where noise and perturbations are added to the data to improve model robustness and performance (Rommel et al., 2022). But the critical difference is that data augmentation is performed in a way that breaks the statistical dependence between the noise features and the outcome, which forces high-capacity ML models to better extract the underlying function of interest. The presence of artifacts induced by peripheral signal generators can therefore not be seen as data augmentation as, in their natural state, they can be correlated with the outcome (as shown here for the two example tasks of age and sex prediction). An interesting twist of this observation would be to develop a proper augmentation approach that injects (simulated) peripheral artifacts into the EEG in ways that break their statistical association with the outcome, hence, enforcing true decorrelation.

Another objection could be that the predictive signal removed with ICA is not only a pure (physiological) artifact but also contains some genuine brain signal. Given that ICA decompositions and component classification can be imperfect, it is a logical possibility that some brain signal is removed along with artifacts and that this led to the observed drop in performance. This would be compatible with the observation that the removal of segments or channels with *autoreject* does not result in the same drop in performance. If ICA removes a brain source it does remove it from the entire recording whereas the rejection of individual segments with *autoreject* is local and preserves brain signals in the retained segments.

For at least three reasons, we hold that regardless of ICA quality, one has to consider that artifacts are predictive and risk diluting CNS biomarkers: 1) prior knowledge of the change in peripheral and body variables in aging and pathology (Golding et al., 2006; Jongkees and Colzato, 2016; Lage et al., 2020; Lindow et al., 2023; Wilkinson and Nelson, 2021). 2) The sensitivity of machine learning models to pick up even weak and hidden patterns. 3) Similar effects were obtained for auxiliary channels where no ICA was applied (Figure 4). The first point deserves some additional reflection. Certain patient populations may be more likely to move, talk or activate facial muscles during the EEG recording than healthy controls. When focusing on a diagnostic analysis and comparing a group of patients with controls, differences in artifact load between diagnoses can therefore lead to a statistical difference that is not driven by differences in brain activity. For example, eye blinks induce major high-amplitude EEG artifacts (Croft and Barry, 2000) and eye blink rate is driven by central dopaminergic function and systematically reduced or increased by pathologies like Parkinson’s disease or schizophrenia, respectively (Jongkees and Colzato, 2016)). Another recent study reported higher artifact probability and fewer clean data segments in children with Fragile X Syndrome as compared to age-matched controls (Wilkinson and Nelson, 2021).

In sum, even if we risk losing some brain signals after ICA cleaning, the alternative would be to have models that cannot be unequivocally interpreted with regard to underlying brain signals.

### Disentangling brain and body EEG generators

Our benchmarks present clear evidence that the signal components attributed to peripheral body signals are predictive themselves, hence, should not be used for a model that intends to capture brain-specific signals (Figure 2-Figure 4). Would it then be appropriate to address the problem of disentangling brain and body signals with statistical techniques for deconfounding (Zhao et al., 2020)? A recent line of work has started studying machine learning techniques for addressing confounding in neuroscience applications (Chyzhyk et al., 2022; Qu et al., 2021). While this can lead to practically useful methods, there is an important theoretical mismatch with our perspective. In confounding, a noise factor of non-interest affects both the inputs and the outcomes. For example, age (cofounder) affects neuronal activity (input) and vascular function (Tsvetanov et al., 2021), both of which influence the blood-oxygen-level-dependent (BOLD) signal measured with functional magnetic resonance imaging (fMRI, outcome). This could induce spurious correlations between electrophysiological measures of neuronal activity and the fMRI signal, as the effects of age on both neuronal activity and vascular function can be mistakenly attributed to a direct relationship between neuronal activity and the BOLD signal. In our theoretical framework, conditions related to the outcome affect latent factors which are mixed in the input signals.

A latent factor view of the problem, therefore, lends itself to trying to disentangle the CNS and peripheral signal generators through blind source separation, ICA and related techniques (Hyvärinen et al., 2004). This is anything but new from the view of traditional EEG analysis, where ICA is a standard tool for isolating brain activity in clinical biomarker studies (Jung et al., 2000). However, this thinking has not yet been broadly embraced in applied ML work with EEG. In this work, we applied the generative model and theoretical results developed in Sabbagh et al. 2019 to better connect the two fields. We reformulated the generative model as a latent factor model with three components: The predictive brain sources, the predictive body sources and the non-predictive noise sources (eqs. 1 to 3). This has motivated signal-isolating regression methods (eqs. eqs. 12 to 15). Our work explored two practical ad-hoc methods for isolating brain from body (Figure 3-Figure 4) for virtually any machine learning pipeline, following a same-analysis approach (Görgen et al., 2018). The implementation of both approaches (ICA versus auxiliary channels) within our wavelet framework has allowed us to gauge direct evidence for the quality of the approximation by comparing power spectra from the respective subspace approximations. With ICA, alpha band oscillations (8-12 Hz) were convincingly isolated and did not seem to leak into the subspace of peripheral models (Figure 3a). On the other hand, alpha band rhythms were clearly visible on the auxiliary channels (Figure 4a). It is therefore unsurprising that the second approach resulted in higher performance estimates for non-brain components as it actually contained some brain signal. The bigger picture was in both cases the same: Mixing of peripheral and CNS signal generators was pervasive across prediction tasks and datasets and deserves explicit handling if the goal is to develop CNS-specific prediction models.

### Wavelets as flexible method bridging predictive modeling and classical analyses

By estimating covariance matrices from convolutions with Morlet wavelets, our framework successfully bridged EEG frequency spectrum descriptors with statistically consistent regression algorithms based on spatial filtering and Riemannian geometry (Barachant et al., 2010; Dähne et al., 2014; Fruehwirt et al., 2017; Gross et al., 2001). These algorithms benefit from theoretical guarantees of zero approximation error under the assumptions (Sabbagh et al., 2020) of constant linear source mixing and a specified nonlinearity (logarithm). Our benchmarks showed that Morlet wavelets can be used as a drop-in-replacement for classical bandpass filtering to obtain the covariance matrix inputs to these types of algorithms without obvious disadvantages, in fact, even leading to improved cross-validation results (Figure 1 – Figure supplement 1).

Overall, our modeling benchmarks replicate the bigger picture reported in previous work (Sabbagh et al., 2020, 2019): The upper triangular vectorization that, essentially, uses the covariances as they are, led to the lowest scores. Of note, this model is statistically consistent if one does not make the assumption of lognormal relationships (Buzsáki and Mizuseki, 2014), leading to linear regression on EEG powers. This is interesting as it was the only method for which our logarithmically scaled wavelets did not consistently lead to improvements over the bandpass filtered equivalents. Models with logarithmic nonlinearity performed consistently better and the *Riemann* model achieved the best results, which taken together suggests that the log-nonlinearity yields models that better match the data generating processes. We noticed that both the theoretically inconsistent *log diagonal* model (biased in the presence of linear source mixing)— representing the classical EEG approach— and the consistent *SPoC* model (Dähne et al., 2014) showed improved performance in our work compared to previously reported benchmark results (Figure 1 – Figure supplement 1), especially if artifact removal was applied (Figure 2). This can be explained in at least two ways. First, we harmonized the processing of covariances across the different pipelines, in particular the handling of data rank and regularization parameters (see methods for details). This was particularly important to enable prediction with minimally processed data that suffer from high-amplitude artifacts, which can lead to ill-conditioned covariance matrices. Second, the improvement could be attributed to the higher performance achieved with the wavelet approach observed across all models (Figure 1 – Figure supplement 1), which could potentially be due to the more adapted frequency smoothing. It is also conceivable that for the less complex *log diagonal* model, the availability of additional frequencies increased model capacity. Compared to previous work (Sabbagh et al., 2020, 2019), where the *SPoC* approach was interpreted as superior to the *log diagonal* approach, here, we observed similar performance for the two models. From the perspective of biomarker development, this would be a welcome result. It gives empirical justification to the less theoretically grounded but far simpler practice in clinical biomarker studies to directly model outcomes from EEG power spectra. Importantly, we do not necessarily expect this result to hold for the (cryogenic) MEG context where the issue of source mixing might be different due to the increased distance between generators and sensors and variable head position. However, the same reasoning might apply for optical MEG (Brookes et al., 2022; Hill et al., 2020).

The *Riemann* model was particularly advantageous when preprocessing was *minimal*, confirming its potential role as an ad-hoc model in early exploratory phases of research as was initially proposed in (Sabbagh et al., 2020). On the other hand, handling this model with intensified data processing was cumbersome as the model makes the strict assumption that the data is full rank (see prediction algorithms - *Riemann* in methods). *SPoC* and the *log diagonal* model, therefore, emerged as potential alternatives as they do not make these strong assumptions.

The more expressive *ShallowNet* (Schirrmeister et al., 2017) did not achieve consistently better results and its performance in the age prediction benchmark (Engemann et al., 2022) was now also reached by the *Riemann* model with wavelets (Figure 1 – Figure supplement 1). In a previous benchmarking study on age prediction, the *ShallowNet* showed clearer advantages over *Riemann* models with a classical filterbank approachk (Engemann et al., 2022). It appears that using a pre-defined Morlet wavelet family to extract temporal features can outperform temporal convolution layers learned in the *ShallowNet* architecture, which, in principle, can overcome the limited fixed-frequency sinusoidal oscillations stipulated by Morlet wavelets. However, it may also simply be a matter of the size of the training data and it does not follow from this that the oscillatory model implied by wavelets is neurobiologically more precise (Cole and Voytek, 2017; Jackson et al., 2019; Schaworonkow and Nikulin, 2019). More importantly, the utility of deep learning is not captured exhaustively by looking at prediction performance in standard settings. As far as custom loss functions, generalization across datasets, or multi-task and multimodal learning are concerned, a deep learning approach is more amenable to the implementation of the latest developments in machine learning research (Banville et al., 2020; Rommel et al., 2022; Wilson et al., 2022). We therefore recommend keeping a good deep learning baseline among the benchmarks in future work.

In sum, machine learning based on covariances derived with log-parametrized Morlet wavelets (as compared to conventional frequency bands) led to improved prediction performance, especially for simpler, hence, potentially more interpretable models. Thus, wavelets emerged as a practical tool to extend established spectral EEG analysis with elements of machine learning.

### A new benchmark for sex prediction from EEG

Interestingly, we observed highly similar trends for sex prediction for both TUAB and TDBRAIN. For sex prediction, only few EEG-based ML studies are available at this point (Jochmann et al., 2023; Khayretdinova et al., 2023; van Putten et al., 2018) and it was a priori not clear if previous methods studied for age prediction would generalize. Comparing our results with recent sex-prediction studies (Jochmann et al., 2023; Khayretdinova et al., 2023) on the TDBRAIN and TUAB datasets show a favorable picture for the models benchmarked in our work.

On the TDBRAIN dataset, Khayretdinova, Zakharov and colleagues (2023) reported a balanced accuracy score of around 80% (± 4.2%) and 84% (± 4.3%) dataset for a gradient boosting and a DCNN convolutional network model, respectively. In this work, scores ranged between 85% and 75% for the best models (± 3% to 5%), depending on the processing applied (Table S1). Precise comparisons are difficult as the authors did not use ICA but rather rejected contaminated segments and no benchmark on minimally processed EEG was provided.

Using a simple amplitude-based convolutional neural network Jochmann and colleagues reported a balanced accuracy score of around 78% (± 2%) on the TUAB dataset with no artifact cleaning. In this work, the *ShallowNet* and *Riemann* benchmarks reached a score around 84% (Table S2) with basic artifact removal and around 80% without artifact removal (±2-5%). Of note, Jochmann et al. 2023 also observed decreases in performance upon applying ICA cleaning (Table 2 in their work).

**Table 2.** IPEG frequency bands.

| Name | $\delta$ | $\theta$ | $\alpha_1$ | $\alpha_2$ | $\beta_1$ | $\beta_2$ | $\beta_3$ | $\gamma$ |
| --- | --- | --- | --- | --- | --- | --- | --- | --- |
| Hz | 1-6 | 6.5-8.5 | 8.5-10.5 | 10.5-12.5 | 12.5-18.5 | 18.5-21 | 21-30 | 30-40 |

Taken together, this argues for the utility of the generative modeling framework and the covariance-based models derived from it beyond its initial exploration for age prediction and brain age (Banville et al., 2023; Mellot et al., 2023; Sabbagh et al., 2020, 2023).

### Additional considerations for model interpretation

Importantly, the proposed framework was not initially motivated by considerations of prediction performance, which emerged as a welcome byproduct of our work. We see the true advantage of the proposed wavelet methodology in its ability to bridge classical results from EEG research on clinical biomarkers with machine learning enabling frequency-by-frequency comparisons between classical measures like EEG power (Figure 1,Figure 3-Figure 4) and machine learning analyses (Figure 3-Figure 5). This not only supports model interpretation but may also facilitate approximations of models by simpler spectral characteristics that can be more easily operationalized and used in clinical studies and subsequent bio-statistical analyses that are highly regulated in drug development and clinical applications.

This brings us to another important point of interest in CNS biomarkers: an interpretation of the signature in terms of brain activity versus variance in brain structure. Our explorations resulted in a mixed picture in which certain frequencies, often around 8-16 Hz, yielded the best results when used as stand-alone models (Figure 5). On the other hand, for most tasks and datasets, above-chance prediction was possible at any frequency and the best performance was obtained by adaptively combining all frequencies instead of averaging them. This signature does not rule out the possibility that the prediction models picked up anatomical differences and ensuing changes in the geometrical configuration of signal generators to the electrodes rather than in brain activity. This hypothesis was considered in related work on sex prediction (Jochmann et al., 2023) and age prediction (Sabbagh et al., 2020) from MEG. In the former work, the researchers noted the predictive importance of spatial patterns and frequencies untypical to EEG. In the latter context, the researchers estimated that around half of the performance might be explainable by source geometry, which was estimated by reconstructing fake covariance matrices from the MEG forward models from individual magnetic resonance imaging (MRI) scans. Alternatively, one might assume that intrinsic long-term autocorrelations are affected by the outcome of interest, translating into changes in the 1/f slope (Cesnaite et al., 2023; Chaoul and Siegel, 2021; Voytek et al., 2015), even if it remains unclear to which extent both explanations might overlap or interact as the link between individual anatomy and brain activity is actively investigated (Pang et al., 2023). While it remains challenging to quantify the impact of anatomical imprinting onto EEG signatures without anatomical measurements, the model inspection techniques presented here can instantly provide an ad-hoc sense of whether the model is driven by specific frequencies or diffuse distributed changes in brain activity that might be related to individual anatomical differences.

As a practical note, the comparisons between broadband covariances and frequency-filtered covariances might be less conclusive in the case of high-density EEG recordings. With five to ten times more channels than with a clinical EEG of 20 channels, additional frequency information may be distinguishable as spatial filtering of EEG implies frequency filtering (Nunez and Srinivasan, 2006). This would be expressed in good performance obtained from broadband covariances in which smaller eigenvectors might encode high-frequency information. Regardless, the proposed methodology will remain of help for exploring the predictive role of specific frequency ranges.

### Limitations and future directions

Our work focused on cross-sectional observational data from two large quasi-public datasets and the two exemplary tasks of sex and age prediction. While this offers versatile opportunities for developing ML models and benchmarking EEG methods relevant for developing biomarkers, we did not predict clinical outcomes in our work. Promising biomarker applications that may be in reach for the methods studied in this work include diagnostic (Blennow et al., 2015), pharmacodynamic (Gautam et al., 2023), prognostic (Lokhande et al., 2022), predictive (Bar-Or et al., 2023; Sechidis et al., 2021) or surrogate-efficacy (Budd Haeberlein et al., 2022; Downing, 2001) questions.

We hope that our work can inform both biomarker developers and machine learning researchers in terms of concepts, methods and empirical benchmarks. We believe that there are several direct applications of our results. Biomarker scientists could reuse our models and techniques on their own clinical data, if the size of the datasets support a machine learning approach. Moreover, the techniques presented here could inform a transfer learning approach where age and sex prediction tasks are used for representation learning (Mourragui et al., 2021) or model predictions are used as downstream variables as in brain age (Cole et al., 2018; Denissen et al., 2022). A second direct application of the models presented here would be proper deconfounding, i.e., when the scientific task requires removing age and sex related components from an outcome of interest (Chyzhyk et al., 2022).

Furthermore, we approximated subspace regression from ICA on individual EEG recordings and automated labeling. The quality of the decomposition therefore stands or falls with the quality of ICA. We hope that our work can inspire the development of novel end-to-end solutions for disentangling brain and non-brain sources, ideally directly built into the prediction models. Promising directions for this effort may lay in the nonlinear ICA (Monti et al., 2020; Zhu et al., 2023), self-supervision (Banville et al., 2020; Tong et al., 2023; Yang et al., 2021) and disentanglement literature (Chen et al., 2018; Lynch et al., 2023; Mathieu et al., 2019; Shu et al., 2018).

Finally, it should be noted that our conclusions are based on two prediction tasks, age and sex. While it is plausible that our findings in principle generalize to other prediction tasks the details (e.g. prediction performance based on peripheral signals, spectral specificity) will be task-dependent.

## Conclusion

Through conceptual analysis, prediction using wavelet-based features and visualization of modeling results across different levels of preprocessing and along the frequency spectrum, our work exposed the risk of applying ML approaches to EEG in the context of biomarker development. Our results emphasize that ML models may not automatically learn the function of interest from mixed signals. When it comes to CNS biomarkers, we think that one has to follow Carl Sagan’s principle that extraordinary claims require extraordinary evidence. This certainly does not question the exploratory value of applied ML in neuroscience and there may be situations in which the best prediction is the priority, regardless of its source. We believe that a new generation of ML techniques is urgently needed to support interpretable disentanglement of latent factors alongside larger clinical trial datasets, potentially enhanced through simulations, for ground-truth assessment and ranking of ML methods for biomarker discovery. To support these developments, in a future updated version of this article, we will share the research code and a Python and Matlab implementation of the log-frequency-parametrized Wavelet method (Hipp et al., 2012) as open-source software with the community.

## Materials and Methods

### Datasets

In this work we used two large quasi-public EEG datasets (institutionally controlled access). We selected these datasets because their size is sufficient for conducting machine learning benchmarks and their demographic and biomedical heterogeneity and EEG setup sufficiently resembles clinical studies.

#### TUAB dataset

The archival Temple University Hospital Abnormal (TUAB) dataset (de Diego and Isabel, 2017) contains a subset of recordings from the Temple University Hospital EEG Corpus (Harati et al., 2014; Obeid and Picone, 2016) that have been annotated as normal or abnormal by medical experts. In the present work we used only the normal recordings (*N* = 1363). The number of female and male participants in this subset is 766 (56%) and 597 (44%), respectively. The age of the participants ranged from 0 to 95 years. EEG was recorded with uninstructed resting state and may therefore contain data from eyes-open and eyes-closed conditions. For convenience, we provide the following description of the TUAB dataset, adapted from our previous work (Engemann et al., 2022): EEG data were recorded using Nicolet devices (Natus Medical Inc.) with 24 to 36 electrodes. The 10-5 system (Oostenveld and Praamstra, 2001) was applied for channel placement. All sessions have been recorded with a common average reference(Nunez and Srinivasan, 2006). Sampling rates varied between 250 Hz and 512 Hz. To the best of our knowledge, the settings for hardware filters are not available.

##### Rationale

We chose this EEG dataset as it represents a heterogeneous sample of the general population of patients from the Philadelphia area seeking medical counseling. Furthermore, the dataset has been popular among applied machine learning researchers (Banville et al., 2020; Darvishi-Bayazi et al., 2023; Gemein et al., 2023, 2020; Sabbagh et al., 2020; Wagh and Varatharajah, 2020; Zhu et al., 2023) and therefore provides a point of reference for algorithmic benchmarking.

##### Data curation and preparation

We identified 21 common channels across recordings which comprise the clinically relevant 10-20 configuration and two mastoid electrodes. Because channel numbers were different between recordings, we re-referenced the data to the average across channels. As the order of the channels was variable, we explicitly reordered all channels consistently. For many patients, multiple recordings were available. For simplicity we only considered the first recording. To ensure comparability, we cropped all recordings to a length of 15 minutes, which was the shortest common recording length.

#### TDBRAIN dataset

The two decades brainclinics research archive for insights in neurophysiology (TDBRAIN) dataset (van Dijk et al., 2022) contains resting state EEG data of 1274 individuals from a heterogenous population of psychiatric patients and healthy volunteers. The age of subjects ranged from 5 to 89 years and the number of female and male participants is approximately equal with 620 female (49%) and 654 male (51%) participants. The recordings were acquired with a 26 channel Compumedics Quickcap or ANT-Neuro Waveguard Cap based on the 10-10 system. In addition to the EEG, seven auxiliary channels were recorded: five channels to measure vertical and horizontal eye movements (electrooculogram; EOG), one to measure the electromyogram (EMG) at the right masseter muscle, and one to record the electrocardiogram (ECG) at the cervical bone. All sessions were referenced against the average of the A1 and A2 mastoids. Hardware filters were set to 0.03 Hz and 100 Hz. Signals were acquired with a sampling frequency of 500 Hz.

##### Rationale

This large dataset was only recently opened for public access and covers a clinically heterogeneous population. The data was acquired using research-grade equipment and a single assessment protocol. The dataset comes with a rich set of auxiliary channels that capture peripheral physiological activity. This renders the TDBRAIN dataset an interesting platform for developing machine learning benchmarks and studying the interplay between CNS and peripheral signals.

##### Data curation and preparation

For many patients, multiple recordings were available. For simplicity we only considered the first recording. Contrary to the TUAB dataset, here, EEG was collected under 2-minute eyes-closed and eyes-open conditions. We pooled the entirety of data, ignoring the conditions.

### Preprocessing

To study the impact of environmental interference and peripheral artifacts on model predictions, we systematically varied the depth of EEG preprocessing across three levels ranging from basic numerically stabilizing processing and harmonization (*minimal* processing) over automated bad segment removal (*autoreject*) to full-blown identification and removal of non-brain artifacts (*autoreject & ICA*).

#### Minimal processing

This level comprises cropping to a recording length of 15 minutes (only applies for TUAB), filtering (FIR filter with pass-band from 1-100 Hz), resampling to a sampling frequency of 250 Hz, epoching into 10 second epochs, and average referencing.

#### Autoreject

This preprocessing level comprises all the steps of *minimal* preprocessing and additional removal and repair of high-amplitude data segments and bad channels using the *autoreject* algorithm (Jas et al., 2017). *Autoreject* is designed to identify and interpolate bad segments and channels based on outlier peak-to-peak amplitude ranges. We used *autoreject* with the following hyperparameters. For consensus, we tested 11 values between 0 and 1 in steps of 0.1. (default). For `n_interpolates` we tested {1, 4, 8} which we adapted to our settings of around 20 EEG channels as we did not want to allow rejecting more than half of the EEG channels (the default would have tested 32 instead of 8). To improve computation time, we used an internal cross validation with 5 iterations instead of the default of 10.

#### Autoreject & ICA

This processing level included all previous steps and added artifact removal via independent component analysis (ICA). For the ICA decomposition, we used a fast approximation of the FastICA model (Hyvärinen et al., 2004) offered by the PICARD algorithm (Ablin et al., 2018) as interfaced through MNE-Python (Gramfort et al., 2014). We used the ICLabel algorithm via its Python implementation in the MNE-ICLabel package (Li et al., 2022) for automatic labeling of the components. Following Rodrigues et al. 2021, components were rejected if any of the artifact probabilities reported by ICLabel (but ignoring the “other” class) were larger than the reported “brain” probability. This labling was later used for ICA-subspace regression (see *Approximate subspace regression and classification*).

### Background: Spectral analysis and machine learning for EEG biomarkers Spectral analysis

As noted early on, EEG comprises oscillatory, i.e. band-limited components and spectrally resolved representations are generally considered useful. A historically grown practice is to evaluate EEG signals in specific frequency bands (alpha, beta, gamma,…). No universally agreed definitions of these bands exist, which results in substantial variability across studies and is hampering progress in biomarker development. There are important efforts to standardize band definitions, e.g. by the IPEG (Jobert et al., 2012). However, such frequency bands are descriptive categories by human observers and not an operating principle of the human brain. Specific definitions of frequencies may not be the right choice for all applications. Moreover, by restricting analyses to pre-specified bands it remains elusive if the choice of these frequency ranges was optimal. We think that given current knowledge about EEG and in the context of ML applications, an unbiased spectrally continuous representation that avoids any band definition is the best choice.

A recent body of literature estimated spectral EEG features (power, phase interactions, power envelope correlations) from complex Morlet wavelets (Forsyth et al., 2018; Frohlich et al., 2019; Hawellek et al., 2022; Hipp et al., 2021) with a logarithmic frequency grid and log-linear scaling of spectral smoothness (Hipp et al., 2012). This approach takes into account prior knowledge about lognormal scaling of brain structure and function (Buzsáki and Mizuseki, 2014), leading to fewer and spectrally wider wavelets with increasing frequency and log-frequency integration over orders of magnitudes (octaves) rather than frequencies. We defined the wavelet families using a base-2 logarithmic grid. As a consequence, the spectral resolution is higher at lower frequencies and spectral smoothing is greater at higher frequencies. Importantly, our implementation parametrizes the wavelets based on their center frequency and their spectral standard deviations in octaves and not in the time domain (Cohen, 2019; Tallon-Baudry et al., 1996).

Details of the implementation and configuration choices are provided below (*Generative latent factor model and CNS-biomarker model*). Spectral power is provided in units of *μV* ^2^*/oct*, which corresponds to *μV* ^2^*/* log_2_(*Hz*). The frequency axis is scaled logarithmically but is labeled in Hz for better readability. For analyzing local frequency effects, we defined groups of five wavelets centered around a center frequency *f* and spanning the range of *f* ·2^−0.5^ to *f*· 2^0.5^ Hz, hence, covering one octave. For comparison against classical approaches based on bandpass filtering in frequency bands Figure 1 – Figure supplement 1, we used the band definitions provided by the IPEG (Jobert et al., 2012), see also Table 2.

### Machine Learning

To improve the generality of our study, we focused on state-of-the-art ML methods for EEG that avoid hand-crafted features motivated by specific theories, clinical populations or cognitive processes (Engemann et al., 2022; Gemein et al., 2020; Zhdanov et al., 2020).

Spatial filtering and Riemannian-geometry (Dähne et al., 2014; de Cheveigné and Parra, 2014; Grosse-Wentrup and Buss, 2008; Roijendijk et al., 2016) enable a general approach that can adapt to specific applications. These methods focus on the between-electrodes covariance matrix as input and therefore can even avoid the pre-specified selection of electrodes or frequencies by algorithmically weighting all inputs (Ang et al., 2008; Sabbagh et al., 2020) through the objective function of the prediction model. They are, therefore, well-suited for isolating the overlapping patterns of distinct EEG-signal generators.

Deep learning (DL) approaches such as convolutional neural networks push this reasoning one step further by not only learning predictive combinations of electrodes but also learning relevant temporal filters (which in turn implies resonance to specific frequencies) from EEG signals (Jing et al., 2020; Schirrmeister et al., 2017; Tveit et al., 2023).

Importantly, results from statistical machine learning in EEG (Sabbagh et al., 2020, 2019) allowed us to map our research question (how EEG-based prediction models are affected by non-brain signals) onto formal model hypotheses (see also, Table 1), which we developed below.

**Table 1.** Models and implied signal-generating hypotheses.

| Model | Hypothesis |
| --- | --- |
| *Upper* (eq. 8) | outcome linear in source power regardless of source mixing |
| *log diagonal* (eq. 9) | outcome linear in log of source power in absence of source mixing |
| *SPoC* (eq. 10) | outcome linear in log of source power regardless of source mixing |
| *Riemann* (eq. 11) | outcome linear in log of source power regardless of source mixing |

## Generative latent factor model and CNS-biomarker model

### Generative model of EEG

The brain contains billions of neurons whose synchronization is reflected in EEG signals. Yet, their activity remains hidden to the observer, and by constraints of linear systems, one cannot distinguish between more linearly independent brain sources of EEG activity than one has EEG channels. Instead of using individual brain anatomy conveyed by MRI scans, machine learning techniques for EEG approach isolation of brain sources statistically through the construct of statistical sources, also known as latent factors.

We extend the model described in Sabbagh et al. 2020, which assumes that the EEG signal ***x*** _*i*_ (*t*) ∈ ℝ^*P*^, recorded on the *i*^*th*^ subject with *P* electrodes, results from the mixing of brain sources. To describe the fact that ***x***_*i*_ (*t*) actually results from the mixing of CNS sources and additional peripheral souces, we denote those by 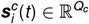 and 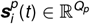 respectively. The resulting generative model of the observed EEG can be expressed as

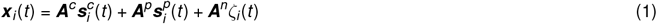

where 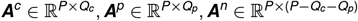 denote the mixing matrices associated with the different signal generators and *ζ*_*i*_ the noise source (not correlated with the outcome). Further assuming that the subspaces of the CNS, peripheral sources and noise are not mixed, we obtain ***A*** = [***A***^*c*^, ***A***^*p*^, ***A***^*n*^] ∈ ℝ^*P×P*^ the mixing matrix and 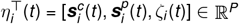 the column vector of sources plus noise, we can then use the compact matrix notation ***x***_*i*_ (*t*) = ***A****η*_*i*_ (*t*).

From this notation, when isolating the CNS and peripheral sources from the noise, we can write 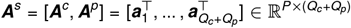, 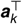 being the *k*^*t*^ *h* column of the mixing matrix and 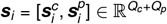 the column vector of sources.

As a result, our statistical modeling can only resolve *P* linearly independent sources, which, obviously, stands in contrast with the complexity of the true biological brain. Of note, this constraint is shared with linear inverse solution techniques such as minimum norm estimates or beamforming (Gross et al., 2001; Hämäläinen and Ilmoniemi, 1994; Van Veen et al., 1997), which project the limited number of linearly independent vectors onto a predefined set of thousands of MRI-defined dipole locations.

### Generative model of outcome and biomarker model

Next we want to formalize outcome measures (including e.g. age, sex, cognitive performance, presence of pathologies, CNS-active drugs) that can modulate source activity and can therefore be predicted from EEG signals.

Denoting ***x***_*i*_ (*t*) ∈ ℝ^*P*^ a time point from the EEG time series recorded using *P* electrodes, and ***X***_*i*_ ∈ ℝ^*P×T*^ the EEG recording of subject *i* with *T* time samples, we can now conceptualize the outcome of interest *y*_*i*_ as a function of the sources *g*(***S***_*i*_) and some additive noise ϵ_*I*_ (Equation 2). A common assumption for *g* is a linear function (weighted sum) of the log of the power of sources ***S***_*i*_ (Sabbagh et al., 2020).

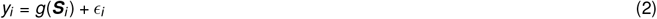

For defining prediction algorithms, it will be convenient to decompose this function into two parts *g*, and *h* which reconstruct the (power of) sources from the EEG signal:

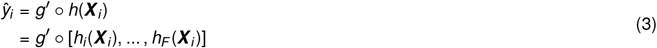

Where:

- 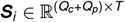: The vector of statistical sources (CNS and peripheral sources).
- 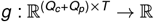 is the true but unknown function generating the outcome *y*_*i*_.
- ***X***_*i*_ ∈ ℝ^*P×T*^ : The EEG recording signal of the subject *i*, with *T* time samples.
- *h*_*f*_ : Function extracting features from the signal in the frequency range defined by the *f* ^*th*^ element of a set of filters that isolate frequencies of interest. This function, and the dimension of the output space will depend on the method used, see prediction algorithms for more details on the different methods. The function is constructed in a way to approximate statistical sources or equipped with invariances to mitigate distorting field spread produced by ***A***^*s*^.
- 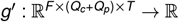 is a function that concatenates the feature vectors and maps them to the outcome. It is based on the estimated representation of the source. This mapping can be a linear function that can be estimated with ridge regression or a more complex nonlinear function.

### Feature Engineering

#### Raw

The *ShallowNet* model (Schirrmeister et al., 2017) operates directly on the epoched (10 second windows) multidimensional time series data (see also section Preprocessing). Here we refer to this as “raw” to indicate that there is no explicit feature extraction step. This is not to be confused with the preprocessing state of the data. Thus, when we refer to the “raw” features, it does not mean that the data has not been preprocessed.

#### Covariance estimation (bandpass filter)

The other baseline models used between-channel covariance matrices estimated from different EEG frequencies as their inputs.

IPEG frequency bands. The classical approach explored in previous work applies bandpass filtering before estimating the covariance matrix (Engemann et al., 2022; Sabbagh et al., 2020). We used the IPEG frequency band definitions designed for pharmacological EEG studies (Jobert et al., 2012).

After bandpass filtering ***X***_*i*_ in frequency *f* we obtain 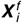. The empirical covariance matrix in that frequency is then given by

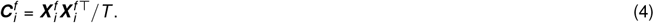

To improve conditioning of the covariance matrices (Engemann and Gramfort, 2015), and for consistency with previous benchmarks, we used Oracle Approximating Shrinkage (OAS) which adaptively downweighs the off-diagonal terms based on the number of samples and number of variables used (Chen et al., 2010). As the covariance was computed over the entire recording, the OAS estimate can be expected to be very close to the empirical covariance.

#### Covariance estimation (Morlet wavelets)

Alternatively, we used convolutions with Morelet wavelets (Morlet et al., 1982) constructed as complex sinusoids windowed by a Gaussian window:

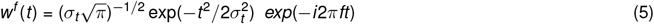

Morlet wavelets have the convenient property that the Gaussian windowing in time, with a standard deviation *σ*_*t*_, translates into a Gaussian smoothing in frequency, with standard deviation *σ*_*f*_ = (2*πσ*_*t*_)^−1^. We used the frequency-domain parametrization described in previous work (Hipp et al., 2012) while implementing a logarithmically spaced grid of frequencies ranging from 1 - 64 Hz. The spectral smoothing was set to *σ*_*f*_ = 0.25 octaves. The spacing of wavelets was set to half the standard deviation, i.e. 0.125 octaves. This resulted in 49 wavelets. To obtain spectral estimates, we convolved the signal ***X***_*i*_ with the complex-valued wavelets *w*^*f*^ (*t*), leading to the complex valued signal ***Z*** ^*f*^ ∈ ℂ^*P×N*^ where *N* is given by the the number of windows and stride length used for numerically approximating the convolution.

The kernel widths were trimmed to 5 standard deviations. For computational efficiency, convolution results were derived at a lower temporal resolution than the original signals, i.e., steps of 1/4 of the kernel width. The covariance was then derived as

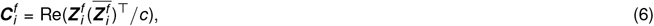

where 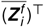 is the conjugate transpose, Re is the real part and *c* is a normalizing constant number representing the number of elements of the convolution result composed of the number valid convolutions defined by the availability of good data segments multiplied by the squared *ℓ*_2_ norm of the wavelet *w*^*f*^ (*t*), which helps estimating the effective sample size *T*. The classical EEG log power-spectral density (PSD) was then derived from the diagonal of the covariance matrix as

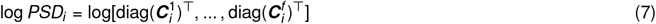

Note that this is identical with the *log diagonal* vectorization used for predictive modeling, which is detailed below.

To improve the conditioning of covariance matrices across different degrees of preprocessing and to ensure that the matrices were positive semidefinite, we applied algorithmic correction to all covariances (Higham, 1988) with a regularization value of 1 × 10^−15^ and covariances scaled in volts squared (*V* ^2^, related to default scaling of the MNE-Python software Gramfort et al. 2013).

### Prediction algorithms

Our benchmark includes four covariance-based methods (*upper, log diagonal, SPoC*, and *Riemann*) and one Neural Network architecture (*ShallowNet*). All of the covariance-based methods consist of a frequency-wise transformation stage followed by a ridge regression (or classification) stage. The regression (or classification) stage is the same across all models such that they only differ in the transformation of the covariance matrix features.

The covariance-based algorithms follow previous work (Sabbagh et al., 2020) and provide useful baselines as they enjoy guarantees under different assumptions about the underlying regression function and degree of signal mixing and can lead to competitive performance in different settings Table 1. Observed differences in prediction performance between these models can, therefore, be practically used to guide interpretation of the underlying regression function and data-generating scenario.

For covariance-based models, we used ridge regression (Hoerl and Kennard, 1970) and ridge classification as supervised learning algorithms. Ridge classification uses ridge regression to predict class labels ∈ {−1, 1 }, such that the decision is obtained from the sign of the prediction. The hyperparameter *α* was controlled through generalized cross-validation (Golub and von Matt, 1997) considering a logarithmic grid of 100 candidate values between 1× 10^−5^ and 1 ×10^10^. This configuration was adapted from previous work (Engemann et al., 2022; Sabbagh et al., 2020). We preferred ridge classification over logistic regression as a probabilistic treatment of predictions was not necessary for this study and hyperparameter selection for ridge classification was fast, hence, well suited for repeated large-scale benchmarking.

#### Upper

The *upper* model (Sabbagh et al., 2020, 2019) vectorizes the upper triangular coefficients of the covariance matrices. This model is thus consistent with a linear relationship between the power and interaction coefficients and the target variable.

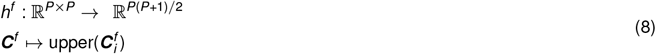

Where *upper* : ℝ^*P×P*^ → ℝ^*P*(*P*+1)*/*2^ is an operator that takes as a vector the upper triangular coefficients of a matrix with off-diagonal terms weighed by a factor 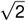 so that the ℓ_2_ norm of the vector is equal to the Frobenius norm of the matrix. Of note, this leads to a *statistically consistent* regression model if no nonlinearity is assumed and the outcome is linear in the source power and not its log (Sabbagh et al., 2020, 2019).

#### log diagonal

The *log diagonal* model (Sabbagh et al., 2020, 2019) extracts the diagonal elements of the covariance matrices (corresponding to the average signal power of each channel) and applies a log transform. This is consistent with a logarithmic relationship between signal power and the target variable. Of note, this leads to a *sstatistically inconsistent* regression model if linear mixing is applied (Sabbagh et al., 2020, 2019).

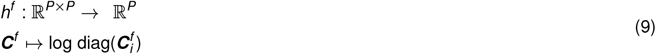

#### SPoC

The Source Power Comodulation (*SPoC*) model (Dähne et al., 2014) aims to learn spatial filters in order to unmix the signal into components with high correlation to the target variable and can be thought of as a regression version of Common Spatial Patterns (Koles et al., 1990). This is similar to Blind Source Separation (BSS) approaches like ICA, where the signal is mapped from sensor space to source space, but special in the sense that the target variable is directly used in the optimization process to maximize correlations between the resulting components and the target variable. This leads to a *statistically consistent* regression model if the outcome is assumed to be linear in the log of the source power.

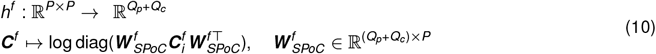

Where 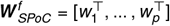 is obtained from solving the generalized eigenvalue problem 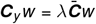 to find the filters that maximize the ratio 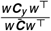 with ***C***_*y*_ denoting the covariances averaged weighted by the outcome and 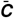 the arithmetic mean of all covariances (Dähne et al., 2014).

#### Riemann

As an alternative to *SPoC*, the *Riemann* model projects the covariances to a Riemannian embedding space, motivated by their positive definite nature (Barachant et al., 2010; Congedo et al., 2017; Sabbagh et al., 2019). In previous work, this has been observed to yield high robustness to noise and good model performance even with minimally preprocessed data. This leads to a *statistically consistent* regression model if the outcome is assumed to be linear in the log of the source power.

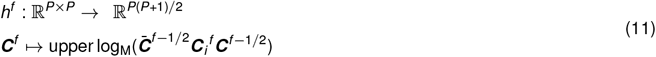

Notably, the *Riemann* model assumes full-rank inputs. In particular, when rank-reducing preprocessing approaches such as ICA are employed, one should therefore be careful not to violate this assumption. Here we ensured that the input has full rank after ICA preprocessing by applying Principal Component Analysis (PCA) as suggested by Sabbagh et al. 2019 by choosing the smallest common rank value, which we obtained from analysis of the eigenvalues of individual covariances.

Of note, it is useful to study both *SPoC* and *Riemann* models as – despite expressing the same signal-generating hypothesis – can behave differently in the face of model violations and noise (Sabbagh et al., 2020, 2019). The *Riemann* model tends to be more robust in the face of noise and model violations. On the other hand, the *SPoC* model is computationally lighter and readily provides compact visualizations of its spatial patterns (Dähne et al., 2014; Mellot et al., 2023), which, taken together, facilitates model interpretation.

#### ShallowNet

The *ShallowNet* architecture proposed in (Schirrmeister et al., 2017) is a convolutional neural network architecture inspired by the Filter Bank Common Spatial Patterns (FBCSP) algorithm (Ang et al., 2008), which can be seen as the classification version of *SPoC*. The architecture presented by Schirrmeister et al. 2017 consists of the following operations

1. Temporal convolution
2. Spatial convolution
3. Square
4. Mean pooling
5. Logarithm
6. Linear regression / classification

and directly maps the raw signal ***X***_*i*_ ∈ ℝ^*P×T*^ to the outcome. The parameters of each layer are described in the original publication. We adapted the *ShallowNet* as a regression model based on a previous publication (Engemann et al., 2022).

Of note, while a proof has never been formally undertaken to the best of our knowledge, the *ShallowNet* expresses the same types of operations as the *SPoC* model. We can therefore assume that the *ShallowNet* can learn the same regression function as the *SPoC* model and can, therefore be *statistically consistent* for the same scenario. As the model has more trainable parameters and can learn the temporal convolution filters, its expressive capacity is higher, hence, it can cover additional regression functions that the *SPoC* model cannot capture.

### Approximate subspace regression and classification

When exploring the relative contribution of peripheral non-brain signals the previous models are used with alternative data inputs to the different feature extractors *h*^(^*f*). These are obtained by applying data selection or processing so that the resulting input can be interpretable as an approximation of the subspaces of interest. In fact, applying the theoretical results from (Absil et al., 2009; Sabbagh et al., 2020), if predictive latent factors related to peripheral body signals are fully silenced through low-rank projection, the remaining subspace leads to a consistent prediction model of brain activity predicting from covariances.

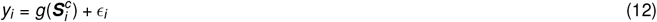

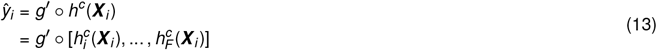

Of note, *h*^*c*^ is a function that approximates reconstruction of the brain sources. This can be inverted, and the same reasoning can be used to define non-brain source models to estimate the relationship between peripheral signals and the outcome.

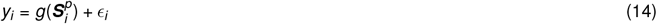

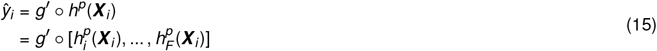

Here, *h*^*p*^ is a function that approximates reconstruction of the peripheral sources. We used two different approaches to estimate the function *h*^*p*^ and to assess the predictive value of physiological (non-brain) signals: (1) the signal reconstructions from ICA artifact subspaces and (2) additional auxiliary channels that were recorded along with the EEG in the TDBRAIN dataset.

#### Subspace regression & classification with ICA

The artifact ICA subspaces provide a complementary source of physiological information that can be extracted from any dataset. First, ICA is used to decompose the data into source components. Next, the components are categorized into brain sources and different classes of artifacts. This step can be automated with a tool like ICLabel (Pion-Tonachini et al., 2019). Finally, the signal corresponding to the artifact subspaces can be reconstructed. Applying eqs. 14 and 15, *h*^*p*^, here, involves reconstructing the EEG from the subspace of ICA components that were labeled as artifact.

Compared to the analysis of auxiliary channels, which requires the acquisition of additional information during the recording, this approach is more flexible and allows us to also extract auxiliary information for the TUAB dataset.

#### Subspace regression & classification from auxiliary-channels

As described in the Dataset section, the TDBRAIN dataset contains auxiliary channels in addition to the EEG channels. These channels are designed to record physiological signals including eye movements, (jaw) muscle activity, and cardiac activity. Applying eqs. 14 and 15, *h*^*p*^, here, involves reconstructing selecting the channels that better expose the artifacts, i.e., the 7 auxiliary channels from the TDBRAIN dataset.

### Statistical Analyses

#### Model performance

To estimate and compare generalization performance, we performed 10-fold cross-validation with reshuffling using a fixed random seed. For the TDBRAIN dataset, which comes with a high cardinality of psychiatric descriptors and diagnoses, we stratified the data to approximately equalize the proportion of psychiatric indications across the cross-validation splits. Of note this only supports qualitative comparisons and cannot be readily converted into hypothesis tests as cross-validation splits are not statistically independent. We practically assessed the chance level by all cross-validation splits exceeding an *R*^2^ of 0 as this score quantifies the improvement over the mean predictor based on the training data. On the other hand, negative *R*^2^ scores indicate prediction failure where the mean predictor has errors than the prediction residual of the model.

#### Hypothesis testing

To compare types of features or degrees of processing across models and datasets, we treated the cross-validation estimates for a given model as a random variable. To obtain uncertainty estimates of pairwise differences in cross-validation performance, we conducted bootstrap resampling with 9999 iterations using the percentile method. We further computed an empirical null-distribution through permutation testing with 9999 iterations. For both analyses, we used the bootstrap and `permutation_test` functions from scipy (Virtanen et al., 2020).

#### Software

All analyses were performed using Python 3.9.17. M/EEG data processing, BIDS conversion and subsequent data analysis steps were carried out with the MNE-Python software (v1.5, Gramfort et al. 2013, 2014), the MNE-BIDS package (v0.13, Appelhoff et al. 2019) and Picard (v0.7, Ablin et al. 2018) for an efficient implementation of FastICA. For artifact removal the *autoreject* package (v0.4.2, Jas et al. 2017) and MNE-ICLabel (v0.4, Li et al. 2022) were used. The joblib library (v1.2) was used for parallel processing. For feature computation, the PyRiemann (v0.4, Barachant 2015) and coffeine (v0.3, Sabbagh et al. 2020) libraries were used. Analyses were composed of custom scripts and library functions based on the Scientific Python Stack with NumPy (v1.24.4, Harris et al. 2020), SciPy (v1.9.1, Virtanen et al. 2020), pandas (v.2.0.3, McKinney and Others 2011) and polars (v0.18.15). For classical machine learning, models were implemented using scikit-learn (v1.3.0, Pedregosa et al. 2011). Deep learning was implemented using the PyTorch Paszke et al. 2019) and braindecode Schirrmeister et al. 2017 packages. All visualizations were performed using matplotlib (v3.7.1, Hunter 2007). Figure 1A was created with BioRender.com.

## Author Contributions

**Conceptualization**: D.E., J.F.H., P.B.

**Data curation**: D.E.

**Formal analysis**: D.E., J.P., P.B.

**Investigation**: D.E., P.B.

**Methodology**: D.E., J.F.H., P.B., P.G.

**Project administration**: D.E.

**Software**: D.E., J.F.H., J.P., P.B.

**Supervision**: D.E.

**Validation**: D.E., P.B.

**Visualization**: D.E., P.B.

**Writing—original draft**: D.E., J.P., P.B.

**Writing—review and editing**: D.E., J.F.H, J.P., P.B., P.G.

## Supporting Information

**Figure 1 – Figure supplement 1.**
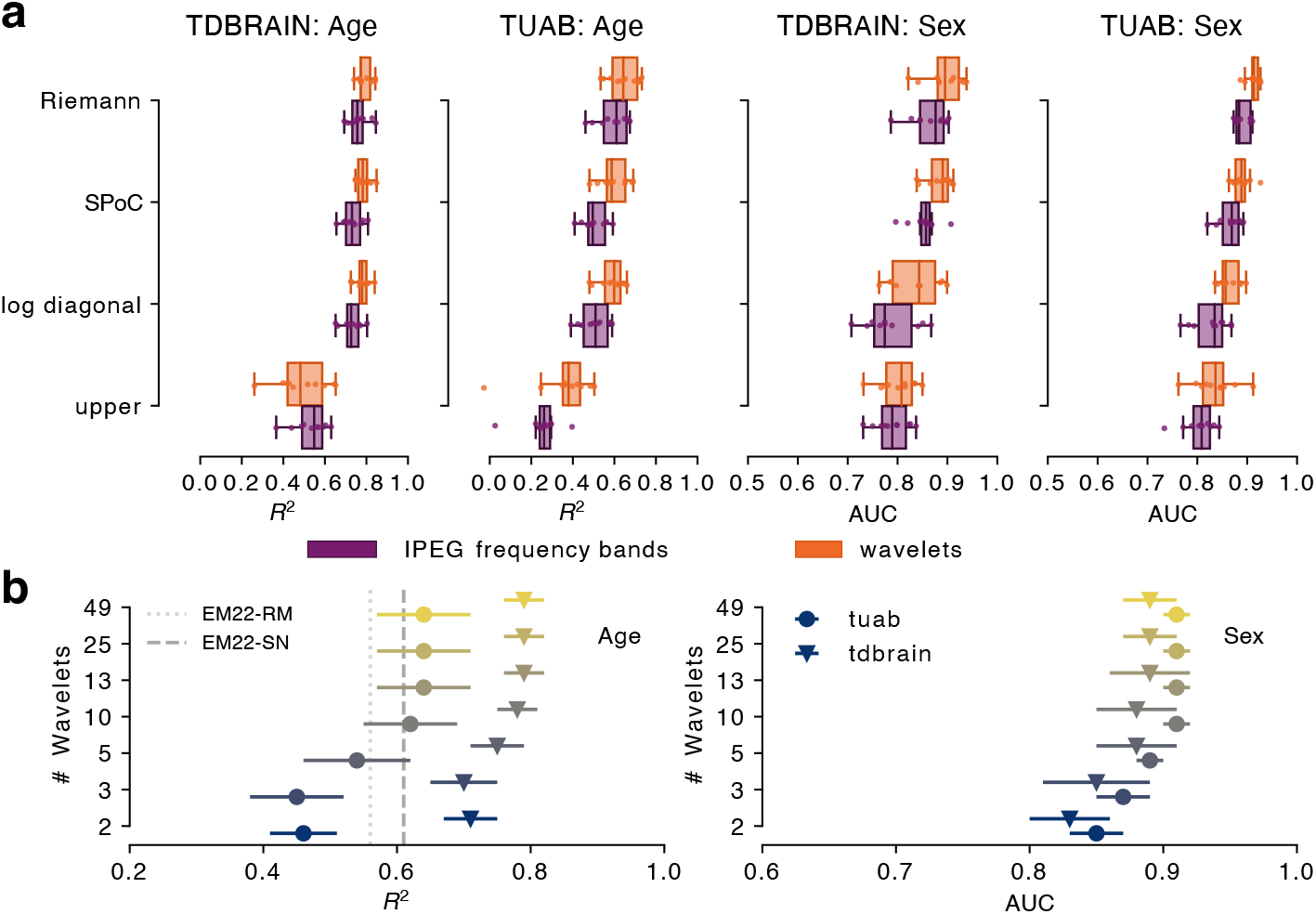
**(a)** Covariance baseline models benchmarked with two alternative sets of covariances along the frequency spectrum. Either the raw signal was bandpass filtered in IPEG (Jobert et al., 2012) frequency bands prior to covariance computation or covariances were computed after convolution with Morlet wavelets. Across models, wavelets consistently led to better results. Wavelets performed on average better by an *R*^2^ of 0.05 (*CI*_95%_ = [0.022, 0.079]) and AUC of 0.03 (*CI*_95%_ = [0.022, 0.036]) across baseline models, tasks and datasets. The *upper* model is an exception as it benefited less consistently. All models with logarithmic nonlinearity visibly improved, which may be related to the explicit logarithmic scaling of wavelets and ensuing balanced representation of signals around the center frequency. For a detailed description of these models,see eqs. 1 to 3 in the main text. **(b)** A high number of wavelets led to favorable performance, saturating above 10 wavelets. EM22-RM and EM22-SN depict the filterbank-riemann and *ShallowNet* benchmarks from (Engemann et al., 2022), respectively. Importantly, the superiority of wavelets over classical frequency bands was, therefore, not trivially driven by the fact that more frequencies were distinguished. Further investigations suggested that the particular selection of wavelets was not critical. Moreover, wavelet-derived covariance features still outperformed the band-pass filtering approach when the wavelets were averaged within the IPEG frequency bands, hence leading to the same number of covariances. This suggests that the distinct factor is not the frequency bands per se but how spectral information is weighted in bandpass filtering as compared to wavelet convolution. Our approach based on (Hipp et al., 2012) integrates by *log*_2_ octaves rather than Hz, hence, providing more equal weighting of low and high frequencies within one bin. Taken together, these results suggest that the established Morlet wavelet methodology can be effectively extended to support covariance-based machine learning approaches and even lead to improvements.

**Figure 2 – Figure supplement 1.**
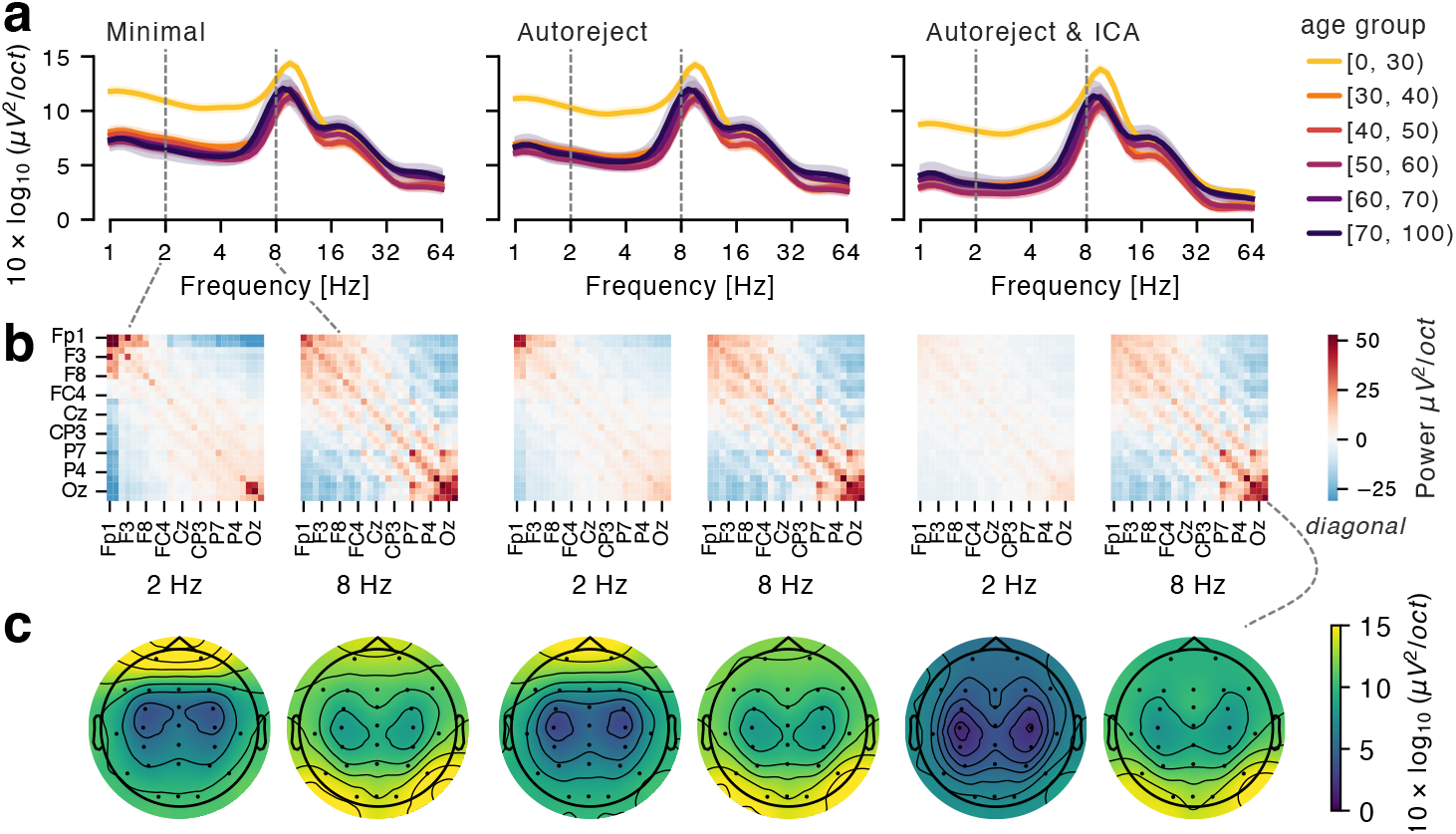
Illustration of spectral signatures across different degrees of preprocessing (*minimal, autoreject, autoreject & ICA*) on the TDBRAIN dataset. **(a)** Power spectral density grouped by age (error bands correspond to 2 standard errors of the mean). Differences in average power spectra across age groups were apparent and dampening of signal power with stronger preprocessing could be observed particularly in low and high frequencies. **(b)** Covariance matrices at 2 and 8 Hz. Activity in neighboring channels was generally positively correlated due to volume conduction. Clusters of high activity emerged in frontal areas at 2 Hz (*minimal, autoreject*) and in occipital areas at 8 Hz. **(c)** Topographic distributions of log power at 2 and 8 Hz, which correspond to the diagonal elements of the covariance matrices. Attenuation of frontal power with increasing preprocessing can be observed, whereas the topography at 8 Hz remains relatively unchanged and reveals the posterior-dominant alpha rhythm. Preprocessing can affect the EEG at different electrode locations and frequencies, which might remove noise or signals that are potentially informative for prediction models based on machine learning.

**Figure 2 – Figure supplement 2.**
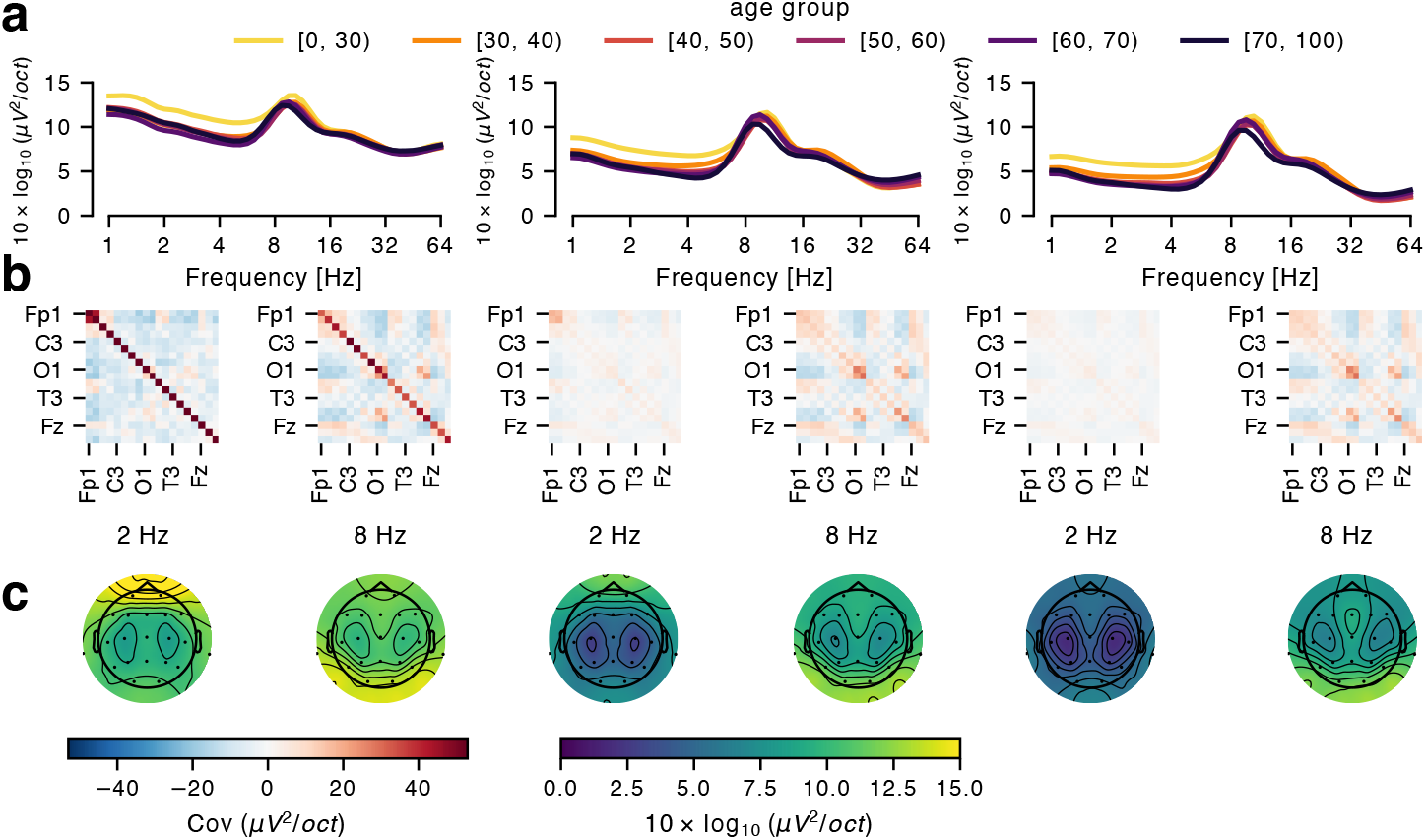
Illustration of spectral signatures across different degrees of preprocessing (*minimal, autoreject, autoreject & ICA*) on the TUAB dataset (same conventions as in Figure 2 – Figure supplement 1).

**Table S1.** TDBRAIN Benchmark Results (10-fold cross-validation).

| Model | Processing | Age Prediction (Regression) |  |  |  | Sex Prediction (Classification) |  |  |  |
| --- | --- | --- | --- | --- | --- | --- | --- | --- | --- |
|  |  | R2 |  | MAE |  | Balanced Accuracy |  | AUC |  |
|  |  | Mean | Std | Mean | Std | Mean | Std | Mean | Std |
| *Riemann* | Minimal | 0.79 | 0.06 | 6.58 | 0.62 | 0.85 | 0.05 | 0.91 | 0.04 |
|  | AR | 0.79 | 0.04 | 6.71 | 0.63 | 0.81 | 0.04 | 0.89 | 0.04 |
|  | AR & ICA | 0.73 | 0.04 | 7.67 | 0.57 | 0.75 | 0.05 | 0.83 | 0.05 |
| *SPoC* | Minimal | 0.75 | 0.06 | 7.2 | 0.79 | 0.81 | 0.04 | 0.89 | 0.02 |
|  | AR | 0.78 | 0.03 | 6.94 | 0.57 | 0.81 | 0.05 | 0.88 | 0.03 |
|  | AR & ICA | 0.75 | 0.05 | 7.41 | 0.85 | 0.77 | 0.04 | 0.84 | 0.04 |
| *log diagonal* | Minimal | 0.77 | 0.04 | 7.09 | 0.69 | 0.78 | 0.03 | 0.85 | 0.04 |
|  | AR | 0.78 | 0.04 | 6.97 | 0.66 | 0.77 | 0.05 | 0.83 | 0.05 |
|  | AR & ICA | 0.76 | 0.04 | 7.22 | 0.77 | 0.73 | 0.05 | 0.8 | 0.05 |
| *Upper* | Minimal | -1249 | 3951 | 34.98 | 64.05 | 0.51 | 0.02 | 0.65 | 0.05 |
|  | AR | 0.49 | 0.12 | 10.48 | 0.8 | 0.72 | 0.03 | 0.8 | 0.04 |
|  | AR & ICA | 0.54 | 0.07 | 10.18 | 0.66 | 0.67 | 0.04 | 0.78 | 0.05 |
| *ShallowNet* | Minimal | 0.64 | 0.13 | 8.21 | 1.95 | 0.75 | 0.04 | 0.84 | 0.04 |
|  | AR | 0.77 | 0.08 | 6.55 | 1.22 | 0.82 | 0.03 | 0.89 | 0.03 |
|  | AR & ICA | 0.75 | 0.08 | 6.83 | 1.09 | 0.76 | 0.03 | 0.84 | 0.03 |

*Note.* Low scores for the *upper* model with *minimal* processing were explained by bad data points leading to bad cross-validation splits. The boxplots in Figure 2 present a robust estimate that is in line with removing bad data points.

**Table S2.**
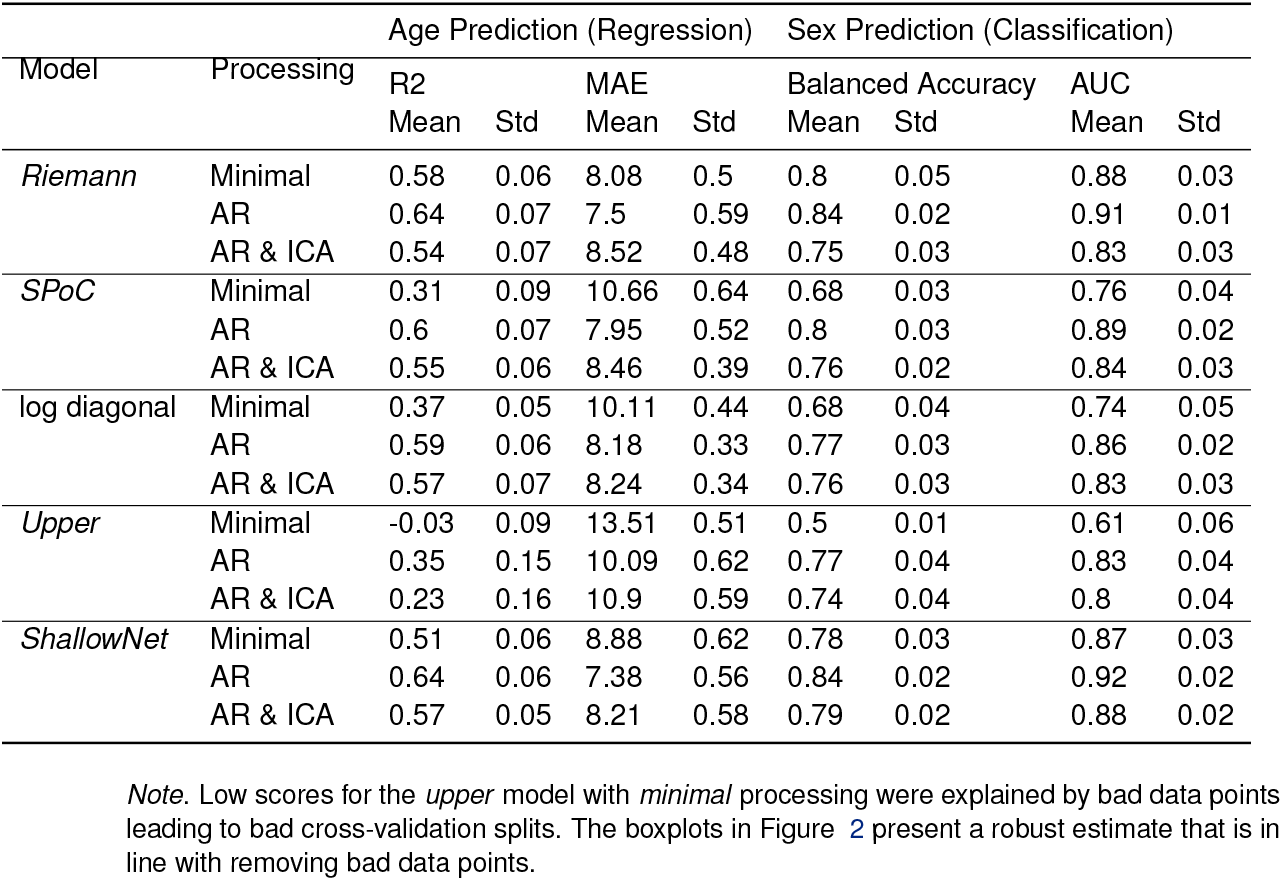
TUAB Benchmark Results (10-fold cross-validation).

